# Rap1 promotes epithelial integrity and cell viability in a growing tissue

**DOI:** 10.1101/2022.10.21.513265

**Authors:** C. Luke Messer, Jocelyn A. McDonald

## Abstract

Having intact epithelial tissues is critical for embryonic development and adult homeostasis. How epithelia respond to damaging insults or tissue growth while still maintaining intercellular connections and barrier integrity during development is poorly understood. The conserved small GTPase Rap1 is critical for establishing cell polarity and regulating cadherin-catenin cell junctions. Here, we identified a new role for Rap1 in maintaining epithelial integrity and tissue shape during *Drosophila* oogenesis. Loss of Rap1 activity disrupted the follicle cell epithelium and the shape of egg chambers during a period of major growth. Rap1 was required for proper E-Cadherin localization in the anterior epithelium and for epithelial cell survival. Both Myo-II and the adherens junction-cytoskeletal linker protein α-Catenin were required for normal egg chamber shape but did not strongly affect cell viability. Blocking the apoptotic cascade failed to rescue the cell shape defects caused by Rap1 inhibition. One consequence of increased cell death caused by Rap1 inhibition was the preferential loss of the polar cells, which later in development led to fewer cells forming a properly migrating border cell cluster. Our results thus indicate dual roles for Rap1 in maintaining epithelia and cell survival in a growing tissue during development.

## Introduction

Epithelia serve critical functions throughout the body’s tissues and organs. For proper homeostasis, epithelia must remain a cohesive unit while being amenable to essential remodeling events. This allows critical epithelial functions such as forming a barrier to pathogens, absorption of nutrients, wound healing, and other important roles (Abramson and Anderson, 2017; Blanpain and Fuchs, 2009; Bröer, 2008; Guillot and Lecuit, 2013; Tai et al., 2019). During development, epithelial tissues undergo dramatic tissue rearrangements while staying intact, for example in convergent extension in *Drosophila* (Irvine and Wieschaus, 1994), ventral enclosure in *C. elegans* (Williams-Masson et al., 1997), and bottle cell invagination in *Xenopus* (Keller, 1981). Once formed, cells in epithelia must maintain polarized cell shapes, stay connected through cell-cell contacts, and survive insults imposed by the tissue environment. Epithelial tissues are challenged by cell turnover, cellular rearrangements, and apoptosis in response to normal tissue growth and homeostasis (Duszyc et al., 2017; Guillot and Lecuit, 2013). Because dysregulation of epithelia shape and cell survival can lead to diseases such as cancer, it is important to understand the mechanisms required for epithelial maintenance.

Here we report a requirement for the conserved small GTPase Rap1 in epithelial maintenance, where it contributes both to cell and tissue shape and to cell viability. The *Drosophila* ovary is an excellent model system to investigate how epithelial cells and tissues respond to challenges such as growth and shape changes during development. The ovary is made up of a series of continuously developing egg chambers. Each egg chamber consists of an inner population of germline derived cells enveloped in a continuous, polarized somatic cell epithelium made of follicle cells. Follicle cells continue to divide until stage 6, when mitosis ceases, resulting in a monolayer of ∼650 cells. The follicle cells undergo a unique rotational migration that helps assemble a basement membrane layer. The basement membrane provides resistance to tissue growth and contributes to egg chamber shape (Duhart et al., 2017). The egg chamber starts out round but eventually grows and elongates to an ellipsoid shape starting in mid-oogenesis. During this process, the inner germline cells expand and press against the follicle cell layer, contributing to the characteristic ovoid shape of the egg. It is critical that the epithelium stays intact to allow successful oogenesis and proper development of a mature, fertilizable egg.

Rap1 plays key roles in epithelial morphogenesis during development, particularly in the establishment of tissue polarity and cell-cell adhesions. In *Drosophila*, Rap1 helps polarize epithelial cells by positioning Bazooka/Par3 (Bonello et al., 2018; Choi et al., 2013) and promotes proper cell-cell adhesion by regulating E-Cadherin-rich adherens junctions (Knox and Brown, 2002). In human podocytes, as well as other cell types, Rap1 regulates integrin mediated adhesion to the basement membrane (Potla et al., 2014). Rap1 also promotes dynamic cell shape changes during development, including the elongation of cells at the leading edge of the lateral epidermis during *Drosophila* dorsal closure (Boettner and Van Aelst, 2007; Boettner et al., 2003).

Here we show that Rap1 GTPase maintains cell and epithelial shapes and promotes follicle cell survival during oogenesis. Loss of Rap1 altered the shape of follicle cells and the egg chamber itself. We find that Rap1 contributes to cell and tissue morphogenesis during mid-oogenesis by ensuring successful actomyosin contractility through the α-Catenin/E-Cadherin adherens junctions in epithelial cells. Notably, inhibition of Rap1 also induced abnormal apoptosis of follicle cells. The pro-survival function of Rap1 was especially important in a specialized pair of epithelial follicle cells, the polar cells, during mid-oogenesis. When Rap1 was inhibited, polar cells failed to maintain Death-associated inhibitor of apoptosis 1 (DIAP1), leading to loss of one or both polar cells. This subsequently led to fewer cells assembling into the migratory border cell cluster. Together, our results reveal dual roles for Rap1 in cell and epithelial morphogenesis and promoting cell viability within a developing tissue.

## Results

### Rap1 GTPase is required for proper egg chamber and epithelial shapes in mid-oogenesis

To better understand the role of Rap1 in epithelial maintenance, we first analyzed the localization of Rap1 using a functional GFP-Rap1 fusion protein expressed under the control of the endogenous *Rap1* promoter (Knox and Brown, 2002). Rap1 is expressed ubiquitously in all cells of the ovary, with highest enrichment at the apical cell cortex of follicle cells, including the polar cells (Fig. 1A). While Rap1 has known roles in the morphogenesis of diverse epithelia (Kim et al., 2022; Knox and Brown, 2002; Wang et al., 2013), few studies have analyzed how Rap1 maintains epithelia during tissue growth. To address this, we inhibited Rap1 activity in the follicle cells using a validated dominant negative Rap1 construct, UAS-DN-Rap1^N17^ (DN-Rap1^N17^), whose expression causes phenotypes that strongly resemble loss of *Rap1* or Rap1 RNAi knockdown phenotypes in the embryo and ovary (Boettner et al., 2003; Perez-Vale et al., 2022; Sawant et al., 2018). DN-Rap1^N17^ was driven by an epithelial-specific GAL4 driver, *c306*-GAL4 (Fig. 1B). Expression of *c306*-GAL4 begins early during oogenesis in anterior and posterior follicle cells, but the highest expression occurs at mid-oogenesis (∼stages 4-8). These stages coincide with a major egg chamber growth phase, that requires the follicular epithelium to stretch and challenges epithelial cell cohesion (Balaji et al., 2019; Crest et al., 2017; Haigo and Bilder, 2011; Spradling 1993).

**Figure 1.**
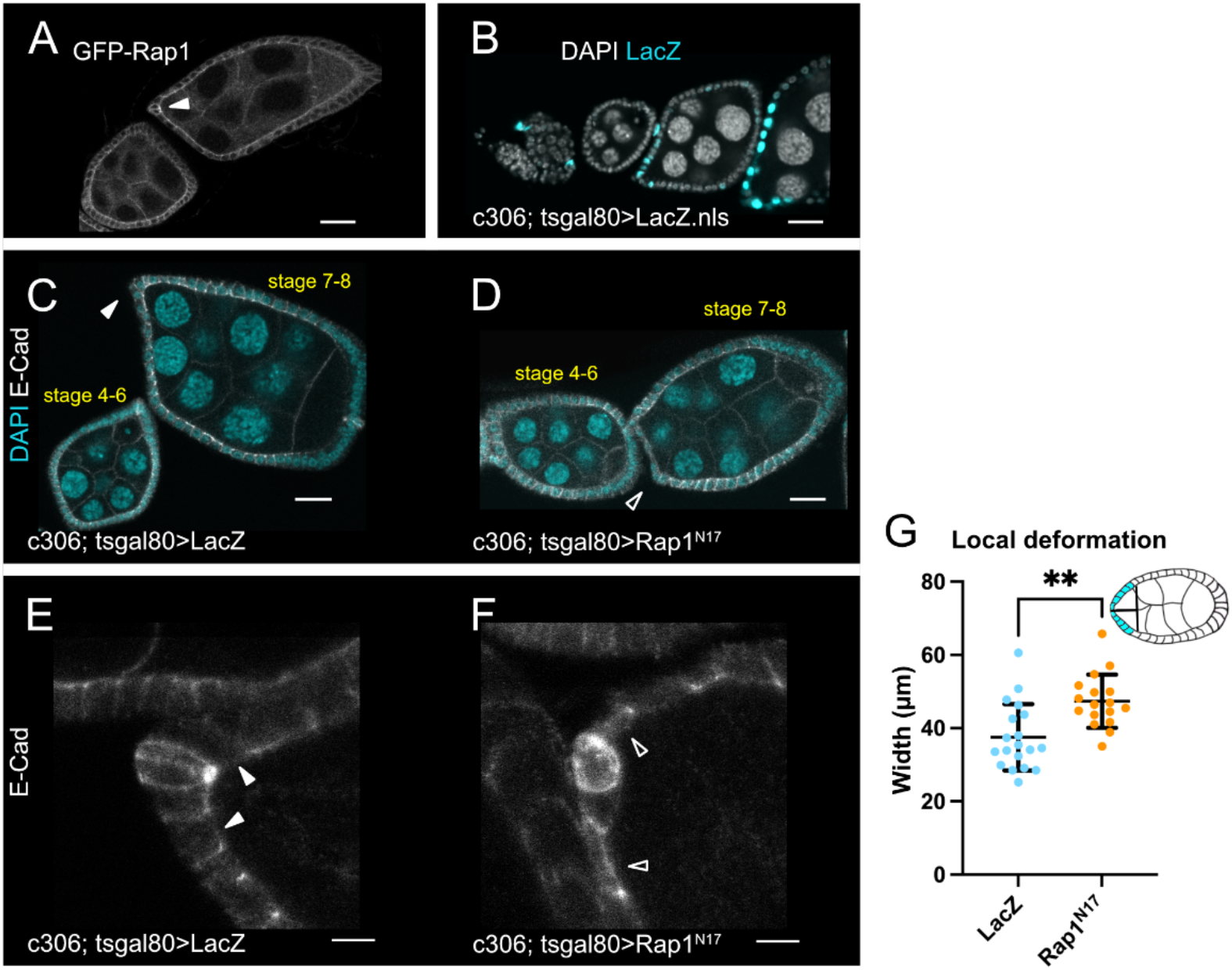
Rap1 is required in the anterior epithelium to maintain follicle cell and egg chamber shapes. (A) Ovariole with GFP tagged Rap1 illustrates Rap1 enrichment in follicle cells. Arrowhead indicates apical Rap1 accumulation in anterior epithelium. (B) Ovariole expressing UAS-LacZ driven by *c306*-GAL4 to demonstrate the *c306*-GAL4 expression pattern. (C-D) LacZ control (C) or DN-Rap1^N17^ (D) egg chambers. Anterior epithelia are distorted at stages 7-8 in DN-Rap1^N17^. Arrowheads indicate anterior region of stage 7-8 egg chambers. Solid arrowhead points to normal anterior epithelium of control (C), while open arrowhead indicates distorted anterior region for DN-Rap1^N17^ (D). (E-F) Anterior regions of stage 7-8 egg chambers showing normal cuboidal epithelium in control (E) or stretched cell shapes in DN-Rap1^N17^ (F). Solid arrowheads indicate cuboidal shaped cells in E. Open arrowheads indicate stretched cells in F. (G) Quantification of local deformation measures taken at 20% of egg chamber length (illustrated by egg chamber schematic in G). ** p≤0.01 two-tailed unpaired t-test. N≥17 egg chambers measured per genotype. (A-D) Scale bars 20μm. (E-F) Scale bars 5μm. Anterior is to the left in this and the following figures.

The shape of egg chambers at stages 4-to-6 appeared to be normal in both control and DN-Rap1^N17^ egg chambers (Fig. 1C-D). However, by stages 7-8, the tissue shape of DN-Rap1^N17^-expressing egg chambers was no longer normal, particularly at the anterior end (Fig. 1D). The anterior of DN-Rap1^N17^ egg chambers was wider and flatter compared to control (Fig. 1C-D). Closer inspection revealed altered anterior epithelial follicle cell shapes specifically in the *c306*-GAL4 expression region (Fig. 1B, D). DN-Rap1^N17^ follicle cells appeared to be stretched, rather than the expected cuboidal follicle cell shapes in control egg chambers at these stages (Fig. 1E-F). We further quantified the observed differences in anterior egg chamber shape, which we termed “local deformation”. To do this, we measured the width of the anterior egg chamber at 20% of the total egg chamber length from the anterior end (*see* Materials and Methods). Control egg chambers retained a characteristic “wedge” shape in this anterior region (mean of 37.49μm). DN-Rap1^N17^ inhibited egg chambers, however, were wider and more “cup” shaped (mean of 47.37μm; Fig. 1G). These data together suggest that Rap1 promotes tissue and epithelial cell shapes during mid-oogenesis.

### Rap1 promotes polar cell shape and apical E-Cadherin accumulation

In addition to the overall distorted anterior follicular epithelial cell shapes in DN-Rap1^N17^ egg chambers, we observed particularly misshapen anterior polar cells compared to control (Fig. 1E-F; 2A-B”). Polar cells are a specialized pair of follicle cells found at each pole of the egg chamber. Polar cells are one of the first specified follicle cells in the ovary and express unique cell markers such as the adhesion protein Fasciclin III (Fas III) (Ruohola et al., 1991). Because anterior egg chamber shape was most impacted by Rap1 inhibition, we focused on the anterior polar cells. We asked when and how polar cell shape was altered by loss of Rap1 activity. To do this, we quantified anterior polar cell shapes by defining their aspect ratio (AR). We measured the width of each polar cell along their dorsoventral (DV) axis and divided by their length along the anterior-posterior (AP) axis (Fig. 2C). Control polar cells at stages 4-6 were typically longer along the AP axis than they were wide along the DV axis (AR=0.61; Fig. 2A, A’, C). At stages 4-6, DN-Rap1^N17^ polar cell shape resembled control (AR=0.65; Fig. 2B, B’, C). Control polar cells maintained their earlier ellipsoid shape at stages 7-8, although they lengthened slightly along the AP axis (AR=0.56; Fig. 2A, A”, C). In contrast, Rap1 inhibited polar cells at stages 7-8 frequently lost this elliptical shape and were now extended along the DV axis, with more spherical shapes (AR=1.02; Fig. 2B, B”, C). Rap1-inhibited egg chambers also had more extreme cases of abnormal polar cell morphology, with either one polar cell that appeared to be missing (Fig. S1A) or a distortion of the polar cell pair, such as pulling apart into “dumbbell” type shapes (Fig. S1B).

**Figure 2.**
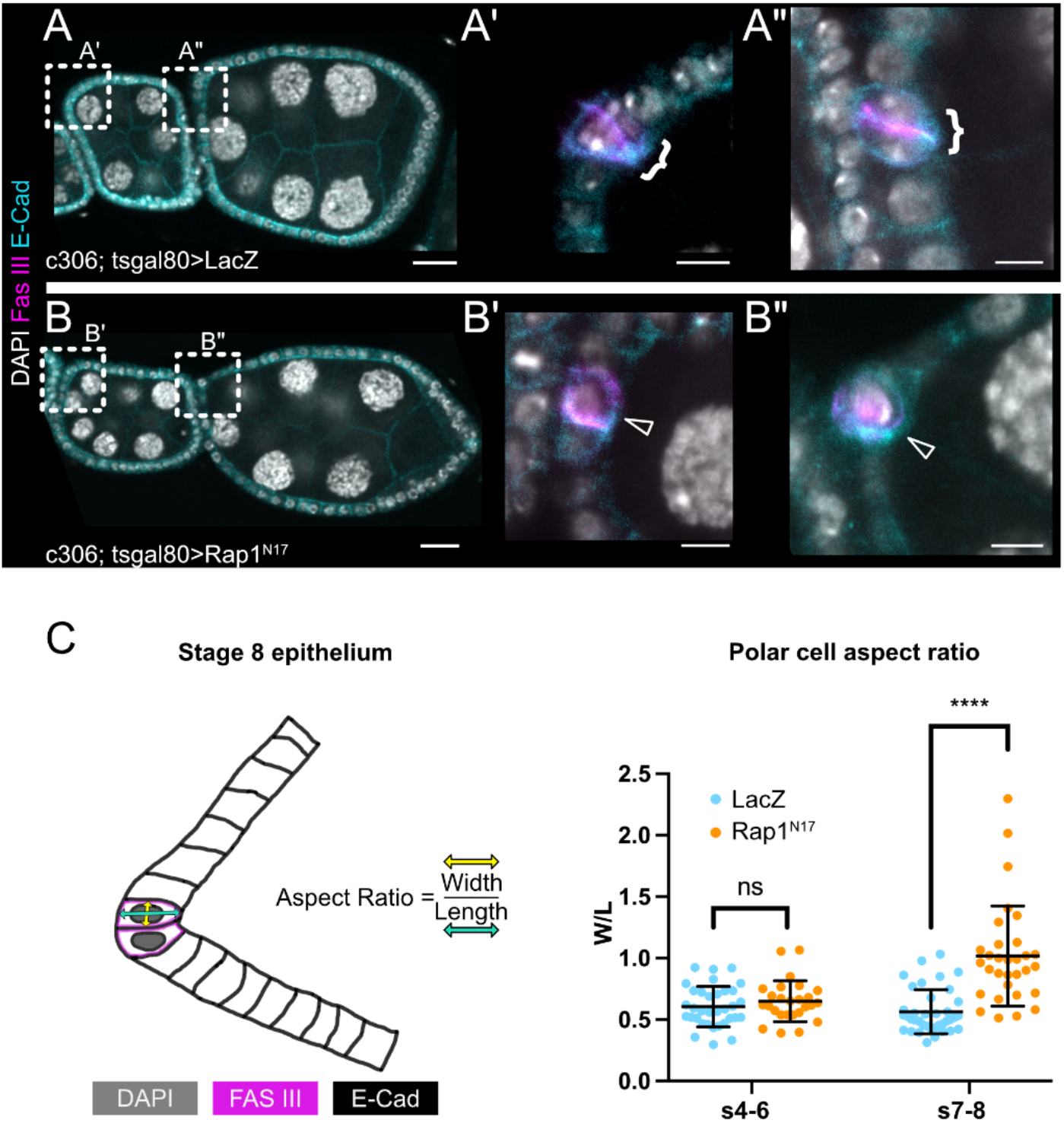
Rap1 is required for proper polar cell shape. (A-B) LacZ (A) and DN-Rap1^N17^ (B) ovarioles imaged for polar cell shape. Dashed boxes indicate the regions expanded in A’, A”, B’, and B”. (A’-A”, B’-B”) Close-up views of anterior egg chamber regions. Control stage 4-6 (A’) and stage 7-8 (A”) egg chamber polar cells. Brackets indicate intact polar cell pairs. (B’-B”) DN-Rap1^N17^ stage 4-6 (B’) and stage 7-8 (B”) egg chamber polar cells. (B’) Only one polar cell is present at stage 4-6. (B”) One spherical polar cell is present at stage 7-8. Open arrowheads indicate only one polar cell is present. (C) Aspect ratio schematic and quantification. Aspect ratio (AR) was defined as the dorsoventral (DV, yellow arrow) width divided by the anterior-posterior (AP, cyan arrow) length. AR measurements plotted for stage 4-6 and stage 7-8 egg chamber polar cells. DN-Rap1^N17^ polar cells are more spherical at stage 7-8 indicated by average AR=1.02 vs average AR=0.56 for control. **** p ≤0.0001 two-tailed unpaired t test. N≥26 polar cells measured per genotype per egg chamber stage. (A-B) Scale bars 20μm. (A’-B”) Scale bars 5μm.

The defective polar cell shapes combined with examples that appeared to split apart prompted us to ask if polar cells in DN-Rap1^N17^ egg chambers were less adhesive. The homophilic cell-cell adhesion protein E-Cadherin accumulates at high levels in polar cells (Niewiadomska et al., 1999). E-Cadherin, as well as the associated cadherin-catenin complex member β-Catenin, highly localizes to the apical side of polar cells, particularly in the region where the polar cell pair is apically constricted (Niewiadomska et al., 1999; Peifer et al., 1993). Therefore, we next investigated if DN-Rap1^N17^ egg chambers accumulated E-Cadherin normally in polar cells (Fig. 3). We measured the apical E-Cadherin fluorescence intensity in stages 7-8 egg chambers along a line drawn starting over the anterior-most nurse cell (“1”; Fig. 3A’, B’) that extended through the apical polar cell-polar cell interface to the lateral interface (“2”; Fig. 3A’, B’, C). In control egg chambers, we observed a peak in fluorescence signal as the line passed through the apical polar cell-polar cell interface (Fig. 3A-A”, C, D). Strikingly, in DN-Rap1^N17^ egg chambers the apical fluorescence enrichment is severely decreased and resembled the intensity values for the portion of the line that extended along the lateral interface (Fig. 3B-B”, C, D). These results indicated that Rap1 controls the localized enrichment of E-Cadherin to the apical interface between polar cells, which in turn may promote the normal shape of the polar cell pair.

**Figure 3.**
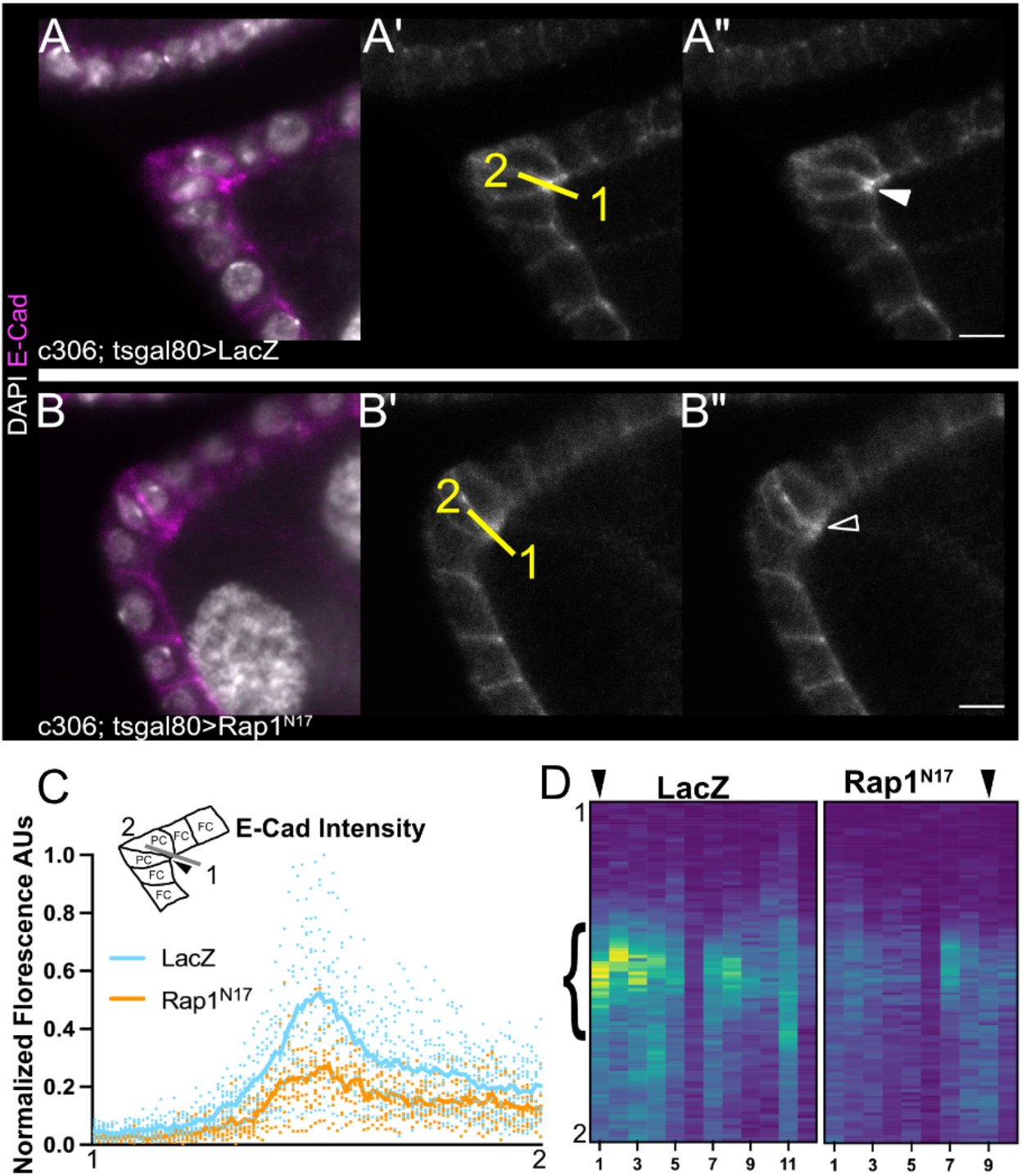
Rap1 is required for apical E-Cadherin enrichment in polar cells. (A-B) Anterior regions of stage 7-8 egg chambers for LacZ control (A) and DN-Rap1^N17^ egg chambers (B). (A’-B”) Single channel panels of A and B showing E-Cadherin (gray) enrichment. (A’, B’) Yellow line indicates region of measurement beginning over nurse cells (1) and ending along lateral polar cell-polar cell interface (2). Solid arrowhead indicates apical accumulation of E-Cadherin. Open arrowhead indicates reduced E-Cadherin enrichment. (A”, B”) Same as A’ and B’ without line overlays for clarity. Image brightness adjusted for presentation purposes. (C) Measurement schematic and quantification of fluorescence intensities measured along lines drawn as in A’ and B’ and normalized to highest signal. Lines represent mean intensity. (D) Heat map representation of C. Bracket indicates area of heat map corresponding to apical enrichment of E-Cadherin in polar cells. Arrowheads at the top of heat map indicate the lanes corresponding to A” and B”. Scale bars 5μm.

Non-muscle myosin II (Myo-II), as visualized with a functionally-tagged regulatory light chain, Spaghetti Squash-GFP, Sqh::GFP (Royou et al., 2004), is enriched across the entire apical surface of the anterior polar cell pair as well as other follicle cells (Fig. S2A-A’’). However, we did not observe any obvious differences in the accumulation of Sqh:GFP at the apical surface of DN-Rap1^N17^ polar cells (Fig. S2B-B”). To confirm this, we quantified Sqh::GFP accumulation by drawing a line across the apical surface of the polar cell pair. We then plotted profiles of the fluorescent intensity values from at least 8 polar cell pairs per genotype. These data indicated no major differences in the Myo-II levels or patterns between control and DN-Rap1^N17^ polar cells (Fig. S2C). Together, our results indicate that Rap1 is dispensable for accumulation of Myo-II but is required for E-Cadherin apical enrichment in polar cells at mid-oogenesis.

### Myo-II and α-Catenin are required for egg chamber morphogenesis

Actomyosin contractility and adhesion both contribute to the shape and integrity of various tissues and organs during development (Harris and Tepass, 2010; Munjal and Lecuit, 2014). Therefore, we wanted to assess whether actomyosin contractility was required for local tissue morphogenesis similar to what we observed for Rap1 (Fig. 4A-E). We targeted the Myo-II regulatory light chain with RNAi-mediated knockdown (Sqh RNAi) and measured local (anterior) tissue deformation. Myo-II deficient staged 7-8 egg chambers resembled what we observed in DN-Rap1^N17^ inhibited egg chambers; the anterior tissue shape was wide and flat (compare Fig. 4B, E to Fig. 1F, G). These results were unexpected given that Rap1 inhibition did not affect apical Myo-II accumulation, at least in polar cells. Therefore, we reasoned that Rap1 may couple actomyosin contractility to the adherens junctions in the anterior epithelium, as has been described in other tissues (Sawyer et al., 2009).

**Figure 4.**
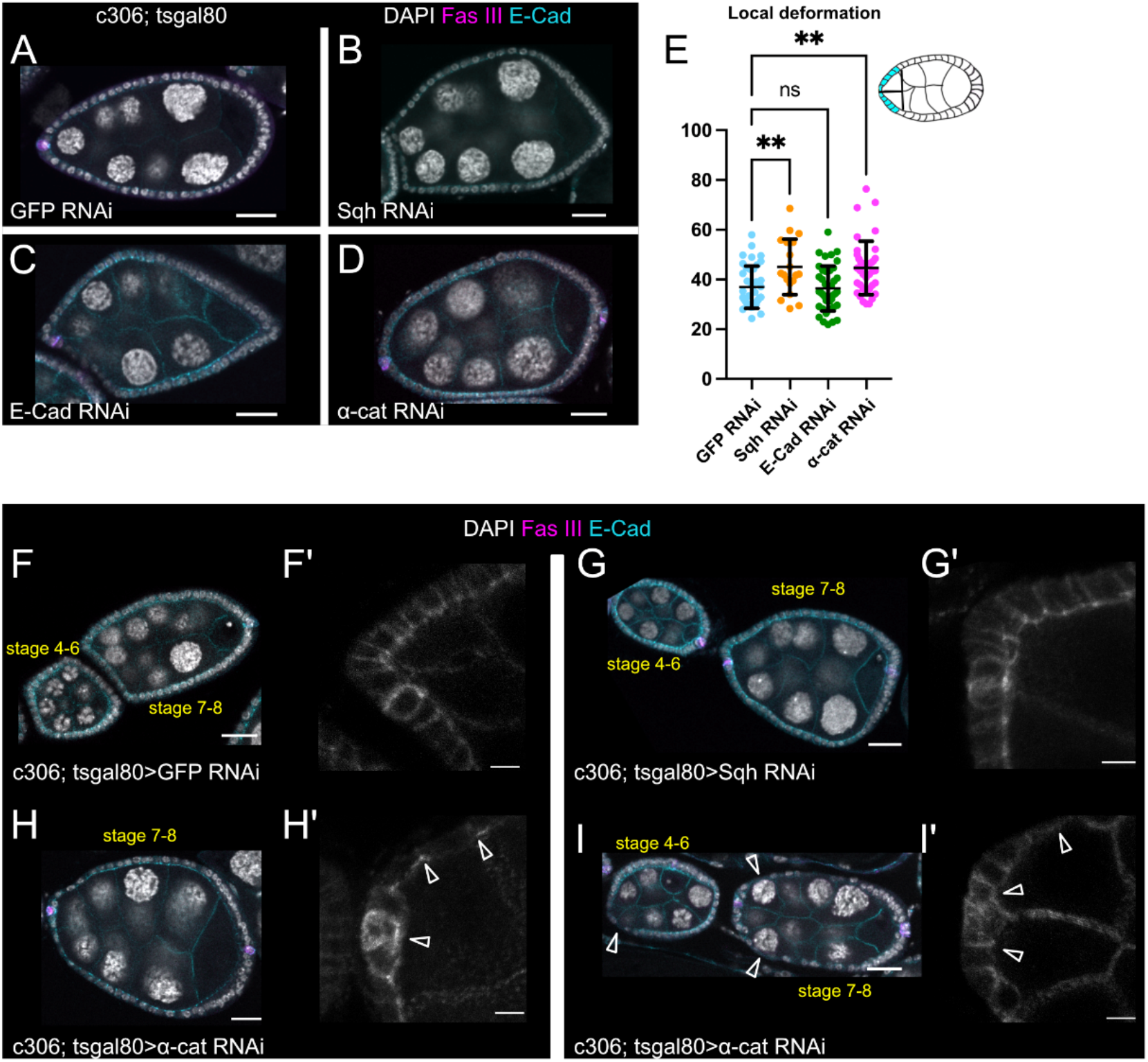
Sqh and α-Catenin maintain local tissue shape and α-Catenin is required for epithelial integrity. (A-D) Stage 7-8 GFP RNAi (A), Sqh RNAi, (B) E-Cadherin RNAi (C), and α-cat RNAi (D) egg chambers imaged for egg chamber shape. (E) Schematic and quantification of local deformation measurement. ** p≤0.01 One-way ANOVA followed by Dunnett’s multiple comparisons test. N≥19 egg chambers per genotype. (F-I) GFP RNAi (F), Sqh RNAi (G), and α-cat RNAi (H-I) ovarioles imaged to show egg chamber shape and anterior epithelium. (F’-I’) Maximum intensity projections of anterior regions of stage 7-8 egg chambers pictured in the main panels. Open arrowheads in H’, I, and I’ indicate “stretched” or clustered cells observed in α-cat RNAi egg chambers. (A-D, F-I) Scale bars 20μm. (F’-I’) Scale bars 5μm.

To test this hypothesis, we next performed RNAi knockdown of E-Cadherin and α-Catenin, members of the adherens junction complex. Surprisingly, E-Cadherin had no effect on egg chamber morphogenesis. Anterior width measurements were no different in E-Cadherin deficient egg chambers when compared to GFP RNAi controls (Fig. 4A, C, E). However, N-cadherin is expressed during these stages and has been shown to undergo compensatory upregulation due to loss of E-Cadherin (Loyer et al., 2015). Thus, we next knocked down α-Catenin (α-Cat), the linker protein that connects cadherin complex members to the cellular F-actin cytoskeleton (Wang et al., 2022; Yonemura et al., 2010). Indeed, downregulation of α-Catenin by RNAi resulted in a wider anterior egg chamber, resembling the tissue shape defects observed with Sqh RNAi and inhibition of Rap1 activity (Fig. 4B, D-E; Fig. 1F-G). Thus, adhesion and actomyosin are required to maintain tissue shapes at mid-oogenesis stages.

We next examined the shape of epithelial follicle cells at the anterior end of the egg chamber when Myo-II or α-Catenin were knocked down by RNAi (Fig. 4F-I’). Close inspection revealed altered individual cell shapes resembling DN-Rap1^N17^ egg chambers only for α-Catenin RNAi (Fig. 4 H-I’) but not Sqh RNAi (Fig. 4G, G’). Notably, we observed a mix of “stretched” or “flattened” follicle cells in α-Catenin RNAi egg chambers along the lateral sides of the egg chamber that looked like the cells in the anterior region of DN-Rap1^N17^ egg chambers (Fig. 4H-I’; compare to Fig. 1F). In the most-affected α-Catenin RNAi egg chambers (Fig. 4I, I’), clumping of cells was observed at the most anterior end and closely resembled the cell shapes reported for *α-Catenin* null mutant follicle cells (Sarpal et al., 2012). The similarities in the tissue and epithelial cell shape phenotypes caused by loss of α-Catenin and Rap1 activity suggested that Rap1 modulates α-Catenin-containing adherens junctions as was observed during dorsal fold formation in the *Drosophila* embryo (Wang et al., 2013).

### Rap1 promotes the viability of epithelial follicle cells during oogenesis

Our analysis of DN-Rap1^N17^ egg chambers at stages 7-8 revealed not only cases of extremely distorted polar cells, but also examples where one polar cell was missing from the required pair (Fig. S1A). These data suggested that Rap1 contributes to cell survival. To test this idea, we first analyzed a marker of apoptosis during oogenesis in control and Rap1 mutant ovaries. We stained egg chambers for antibodies to cleaved death caspase-1 (cDcp-1), which recognizes both *Drosophila* effector caspases, Death caspase-1 (Dcp-1) and Death related ICE-like caspase (Drice) (Li et al., 2019). During early oogenesis, excess polar cells (“supernumerary” polar cells) and excess stalk cells form but are eliminated by apoptosis during stages 3-6, whereas in healthy ovarioles other follicle cells do not undergo cell death (Borensztejn et al., 2013; Borensztejn et al., 2018; Khammari et al., 2011; Lebo and McCall, 2021; cDCp-1+ cells; Fig. 5A). However, in DN-Rap1^N17^ ovarioles, in addition to the normal pattern of early polar cell and stalk cell apoptosis, we observed additional caspase activity in follicle cells during stages 2 through 8 of oogenesis (cDCp-1+ cells; Fig. 5B). We quantified the number of cDcp-1 cells in control and DN-Rap1^N17^ ovarioles and saw a significant increase in apoptotic cells only when Rap1 activity was inhibited (Fig. 5C). Activation of Rap1, through expression of constitutively active CA-Rap1^V12^ did not decrease apoptotic cells during any stage of oogenesis (Fig. 5C). These data suggest that Rap1 is required for follicle cell survival during oogenesis but is not sufficient to block normal developmental apoptosis.

**Figure 5.**
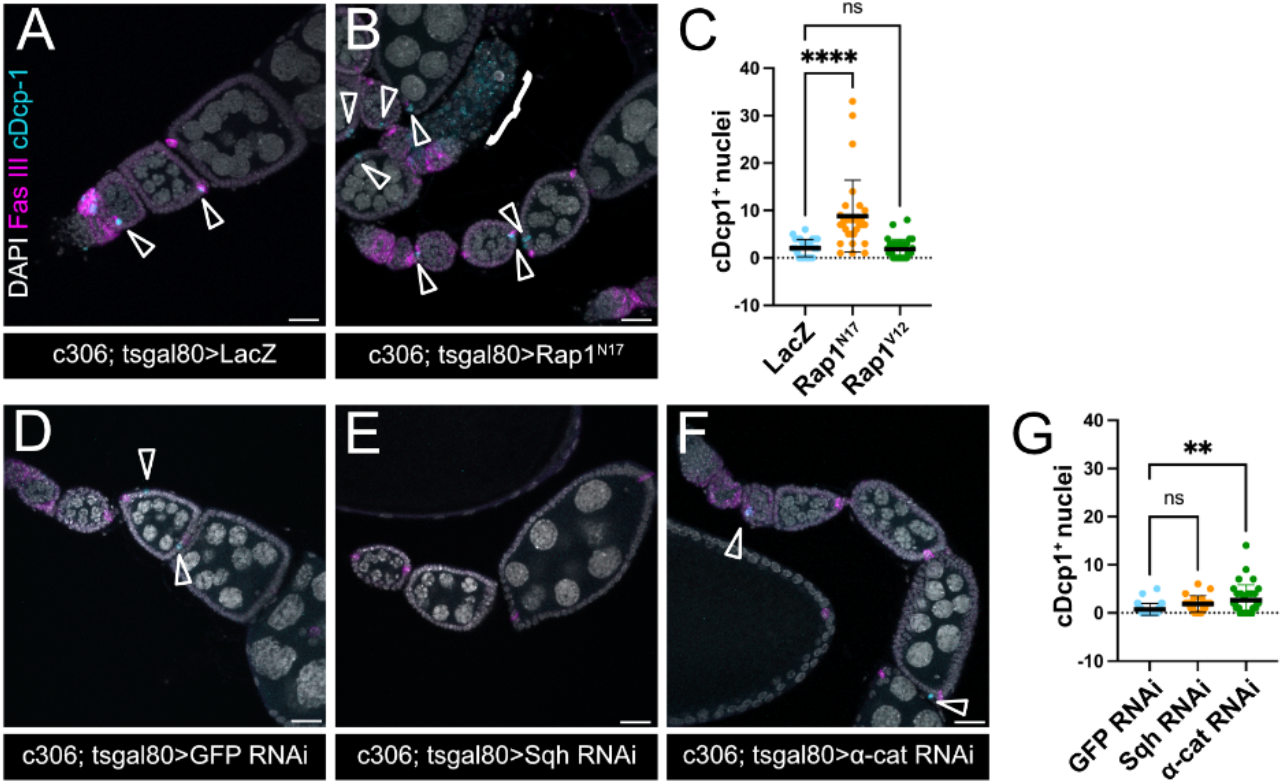
Rap1 and α-Catenin are required for cell viability during oogenesis. (A-B) Maximum intensity projections of ovarioles showing caspase positive nuclei for LacZ (A) and DN-Rap1^N17^ (B). Open arrowheads in A and B indicate cDcp1 positive nuclei and the bracket in B indicates a degenerating egg chamber. (C) Quantification of cDcp1 positive nuclei per ovariole (egg chamber stages 2-8 only). DN-Rap1^N17^ ovarioles have greater number of cDcp1 positive nuclei. **** p≤0.0001 One-way ANOVA followed by Dunnett’s multiple comparisons test. N≥29 ovarioles per genotype. (D-F) Maximum intensity projections of ovarioles for GFP RNAi (D), Sqh RNAi (E), and α-cat RNAi (F). Open arrowheads indicate cDcp1 positive nuclei. (G) Quantification of cDcp1 positive nuclei. ** p≤0.01 One-way ANOVA followed by Dunnett’s multiple comparisons test. N≥20 ovarioles per genotype. Scale bars 20μm.

Because Rap1 promotes both follicle cell survival and epithelial and tissue morphogenesis during oogenesis, we next asked if Myo-II or α-Catenin were also similarly required for follicle cell viability. We analyzed cell death during oogenesis using cDcp-1 in Sqh RNAi, α-Catenin RNAi, and matched control (GFP RNAi) ovarioles. We observed minimal cell death in Sqh RNAi and GFP RNAi controls (Fig. 5D, E, G). However, there was a significant increase in cDcp-1 positive cells for α-Catenin RNAi ovarioles (Fig. 5F, G). These results suggest that while both Myo-II and α-Catenin regulate egg chamber morphogenesis, only α-Catenin is required for viability of the follicle cells. Notably, although α-Catenin RNAi results in an increase in cDcp-1 positive cells, fewer cells per ovariole were cDcp-1 positive than when Rap1 was inhibited (2.6 cDcp-1-positive cells for α-Catenin RNAi, Fig. 5G, compared to 8.8 cDcp-1-positive cells per ovariole for DN-Rap1^N17^, Fig. 5C). These findings suggest that cell adhesion via α-Catenin is required for cell viability, though Rap1 may have a greater role in promoting cell survival.

### Rap1 promotes polar cell survival by suppressing apoptosis, but polar cell shape defects are not apoptosis-dependent

As described above, loss of Rap1 activity causes death of follicle cells and loss of polar cells (Fig. 5B-C; Fig. S1A), in addition to defects in polar cell shape (Fig. 2B-C; Fig. S1B). Therefore, we wanted to determine if Rap1 was required to manage elimination of supernumerary polar cells that form at each pole of early egg chambers (Besse and Pret, 2003; Borensztejn et al., 2013; Khammari et al., 2011). Removal of the extra polar cells by apoptosis ensures that only two “mature” polar cells remain at anterior and posterior egg chamber poles by stages 5/6 (Besse and Pret, 2003; Borensztejn et al., 2013; Khammari et al., 2011). One possibility for the absence of polar cells in later staged DN-Rap1^N17^ egg chambers could be a requirement for Rap1 in early polar cell development and elimination. Alternatively, Rap1 could promote the survival of mature polar cells later in oogenesis after these cells are specified.

We examined anterior and posterior polar cells in egg chambers from stages 3-10 of oogenesis using Eyes Absent (Eya) and FasIII (Fig. S3A). Eya suppresses polar cell fate and is present in all follicle cells except polar cells (Bai and Montell, 2002) and FasIII specifically marks the polar cell pair (Ruohola et al., 1991). We observed similar numbers of supernumerary polar cells at early stages of development (e.g., stages 3-4 and stages 5-6) in control and DN-Rap1^N17^ (Fig. S3A). Using CA-Rap1^V12^, we next asked if increased Rap1 activity resulted in extra polar cells being maintained until maturity. At each stage examined, we observed similar numbers of polar cells for CA-Rap1^V12^ and controls (Fig. S3A). Moreover, in all genotypes, in agreement with other studies, there was an overall decrease in the number of polar cells as oogenesis progressed (Fig. S3A; Borensztejn et al., 2013; Khammari et al., 2011).

Since early polar cell development appeared normal in Rap1-deficient egg chambers, we next asked if Rap1 regulated the survival of mature polar cells later in oogenesis. We tracked polar cell-specific accumulation of cDcp-1. Control “mature” polar cells at stages 7-8 rarely died (Fig. 6A-A”;1 out of 66 egg chambers). In contrast, we observed an increased frequency in the accumulation of cDcp-1 in staged 7-8 DN-Rap1^N17^ polar cells, well after the conclusion of normal developmental apoptosis (Fig. 6B-B”; 7 out of 61 egg chambers). These polar cells were frequently pyknotic as visualized by DAPI staining (Fig. 6B’). These data together indicate that Rap1 promotes the survival of both follicle cells and mature polar cells during oogenesis.

**Figure 6.**
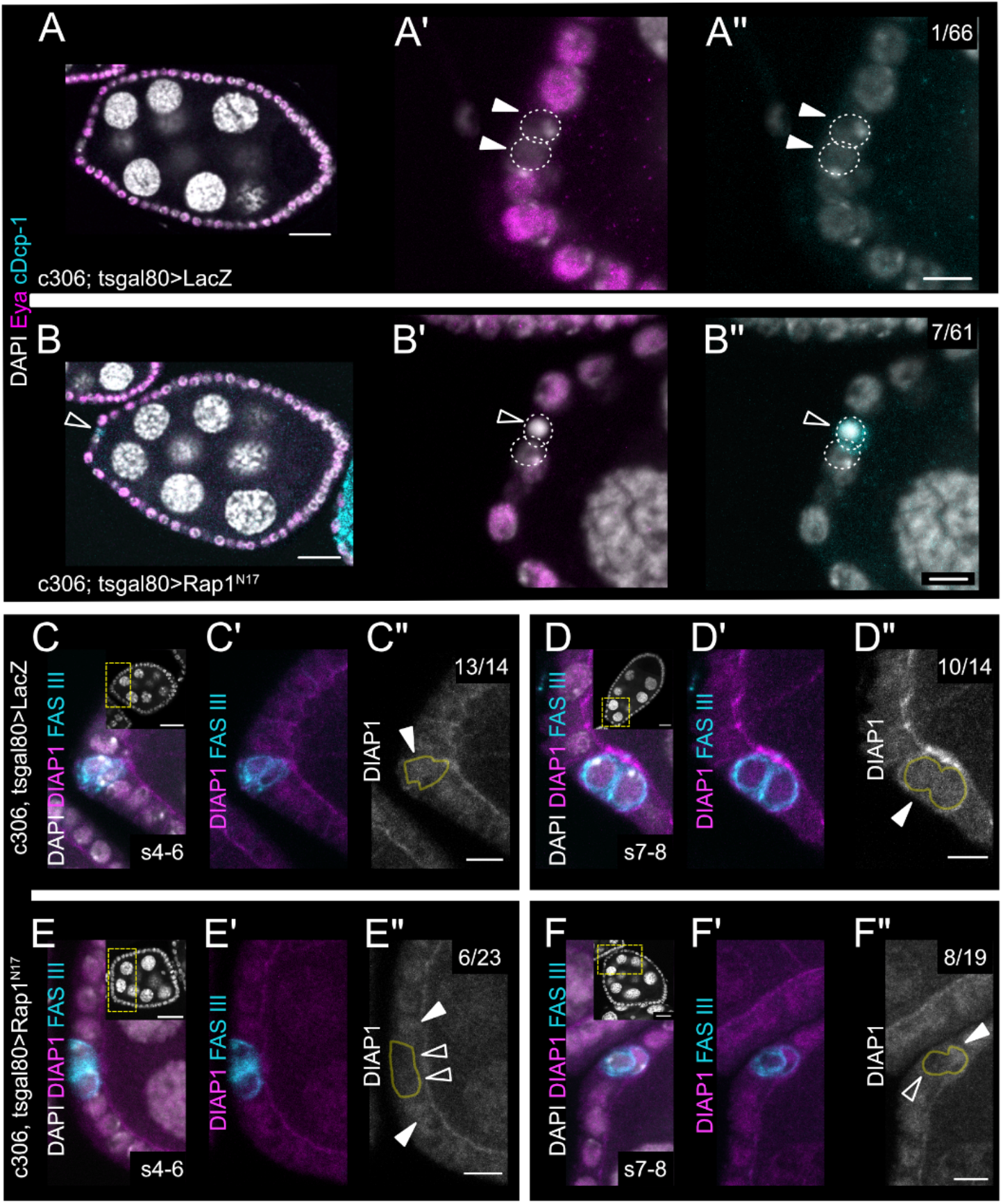
Rap1 is required for polar cell viability and proper DIAP1 accumulation. (A-B) Stage 7-8 LacZ (A) and DN-Rap1^N17^ (B) egg chambers scored for cDcp1 positive polar cells. (A’, B’) Close-up views of anterior regions of the egg chambers depicted in A and B, showing only DAPI and Eya staining. (A”, B”) Same images as A’ and B’ showing only DAPI and cDcp1. Filled arrowheads in A’ and A” indicate viable mature polar cells. Open arrowheads in B’ and B” indicate a cDcp1 positive dying polar cell. Numbers in A” and B” indicate number of observations of cDcp1 positive polar cells per stage 7-8 egg chambers scored. (C-C”, E-E”) Stage 4-6 LacZ (C-C”) and DN-Rap1^N17^ (E-E”) egg chambers. (C’, E’) Same images as C and E showing only DIAP1 and FasIII. (C”, E”) DIAP1 only. (D-D”, F-F”) Stage 7-8 LacZ (D-D”) and DN-Rap1^N17^ (F-F”) egg chambers. (C-F) Insets show DAPI-stained whole egg chambers corresponding to zoomed regions with yellow boxes outlining the approximate region depicted in main panels. (C”-F”) Polar cells are outlined in yellow. Solid arrowheads indicate normal DIAP1 accumulation. Open arrowheads indicate reduced DIAP1. The number of egg chambers with normal DIAP1 accumulation out of total egg chambers scored is reported in the upper right of C”, D”, E”, and F”. (A-F) Scale bars 20μm. (C”-F”) Scale bars 5μm.

To further characterize this increase in polar cell death when Rap1 was inhibited, we next examined the apoptotic cascade. Death-associated inhibitor of apoptosis 1 (DIAP1) blocks caspase activity and thus prevents cells from undergoing apoptosis (Hay et al., 1995; Yan et al., 2004). DIAP1 specifically accumulates in the two mature polar cells and promotes their survival, whereas DIAP1 is downregulated in the supernumerary polar cells leading to their developmental apoptosis (Borensztejn et al., 2013; Khammari et al., 2011). We reasoned that a decrease in DIAP1 might precede the death of mature polar cells observed in DN-Rap1^N17^ egg chambers. Therefore, we analyzed DIAP1 accumulation in polar cells relative to their most adjacent follicle cell neighbors at both stages 4 to 6 and at stages 7 to 8. At the earlier stages (stages 4-6), we observed normal accumulation of DIAP1 in 13 out of 14 control polar cells (Fig. 6C-C”). However, DIAP1 accumulated in only 6 out of 23 DN-Rap1^N17^ polar cells (Fig. 6E-E”).

Similarly, at later stages, DIAP1 accumulation was lower in DN-Rap1^N17^ inhibited polar cells, with only 8 out of 19 egg chambers having normal DIAP1 accumulation (Fig. 6F-F”), compared to 10 out of 14 for control egg chambers (Fig. 6D-D”). Interestingly, DIAP1 levels were altered at a greater frequency in DN-Rap1^N17^ egg chambers (Fig. 6E-F”) than the observed frequency of polar cell death (Fig. 6B”). One possibility is that not all cells that lose DIAP1 accumulation undergo cell death. Alternatively, the threshold for DIAP1 protein depletion may need to be relatively severe for apoptosis to occur. Taken together, our results favor a role for Rap1 in suppressing the apoptotic cascade through regulating the levels of DIAP1 in mature polar cells.

We next asked if the polar cell morphology defects caused by loss of Rap1 activity were due to apoptosis. To test this, we asked if blocking the apoptotic cascade could rescue polar cell shape defects caused by loss of Rap1 activity (Fig. S3B-D). We performed polar cell aspect ratio measurements in egg chambers that co-expressed DN-Rap1^N17^ with either a LacZ control or with the apoptosis inhibitor, baculoviral p35 (Clem et al., 1991; Hay et al., 1994). We found that co-expression of p35 along with DN-Rap1^N17^ failed to rescue the polar cell aspect ratio defects compared to co-expression with LacZ (Fig. S3B-D). There was no statistical difference in the aspect ratios of DN-Rap1^N17^ polar cells either co-expressing the LacZ control or p35 (AR=0.84, DN-Rap1^N17^ + LacZ; AR=0.85, DN-Rap1^N17^ + p35; Fig. S3D). These results provide further evidence that Rap1 likely controls polar cell morphogenesis independently of its function in promoting cell survival.

### Rap1 requirement for polar cell survival supports formation of border cell clusters with optimal cell numbers

The role for Rap1 in promoting polar cell survival prompted us to ask what the developmental consequences were later in oogenesis. During late stage 8, the anterior pair of polar cells specifies which follicle cell neighbors become the migratory border cells through the secretion of the JAK/STAT ligand Unpaired (Beccari et al., 2002; Silver and Montell, 2001). Between 4-8 follicle cells activate high levels of JAK/STAT and subsequently surround the polar cells to produce a border cell cluster with a total of 6 to 10 cells (Silver and Montell, 2001). Having an optimal number of cells helps the border cell cluster efficiently reach the oocyte at the correct time (Cai et al., 2016; Starz-Gaiano et al., 2008; Stonko et al., 2015)

Because inhibition of Rap1 caused a frequent loss of mature polar cells, we determined how this impacted the size of the migratory border cell cluster. We reasoned that losing a polar cell might decrease the number of cells found in border cell clusters. We quantified the total number of cells found within control (UAS-LacZ) versus Rap1-deficient (UAS-Rap1^N17^ or UAS-Rap1 RNAi) border cell clusters (Fig. 7 A-C). When Rap1 was inhibited using with DN-Rap1^N17^ driven by *c306*-GAL4, we found a strong reduction in the average number of cells per cluster compared to control (Fig. 7A’, B’; average of 6.1 cells in control compared to 4.6 cells in DN-Rap1^N17^). *c306*-GAL4-driven *Rap1* RNAi also reduced the number of cells per cluster to an average of 5.2 cells compared to 6.1 cells in *mCherry* RNAi controls (Fig. 7C). We analyzed the number of cells in border cell clusters using a different GAL4, *slbo-*GAL4, which is expressed later than *c306*-GAL4 and in a more restricted pattern (Fig. S4A). The overall number of cells in *slbo-*GAL4 control egg chambers is higher than that observed for *c306-*GAL4 controls (average of 8.2 cells; Fig. S4B, B’, D). Nonetheless, we observed a significant reduction in the number of DN-Rap1^N17^ cells within the border cell cluster (Fig. S4C-D; average 6.7 cells). Thus, Rap1 is required for migrating border cell clusters to have an optimal number of cells.

**Figure 7.**
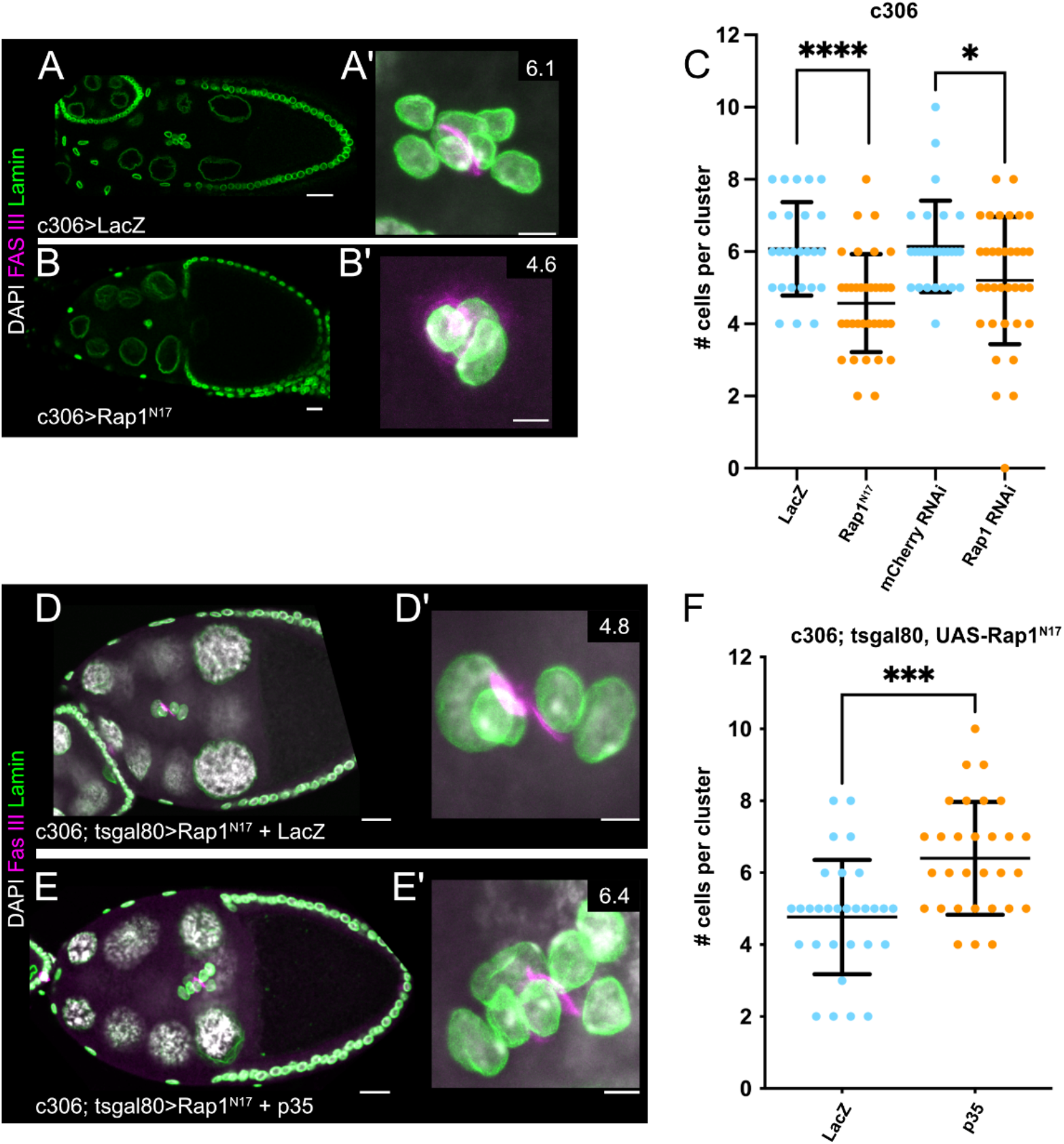
Rap1 dependent cell viability is required for proper border cell cluster assembly. (A-B) Stage 9-10 egg chambers for LacZ (A) and DN-Rap1^N17^ (B). (A’-B’) Maximum intensity projections of border cell clusters from egg chambers pictured in A and B. Numbers at top right are average number of cells observed for each genotype. (C) Quantification of cell number per cluster for each genotype. * p≤0.05 ****p≤0.0001 two-tailed unpaired t test. N≥26 border cell clusters per genotype. (D-E) Stage 9-10 egg chambers for DN-Rap1^N17^ + LacZ expression control (D) or DN-Rap1^N17^ + p35 (E). (D’-E’) Maximum intensity projections of border cell clusters from egg chambers pictured in D and E. Numbers at top right are average number of cells observed for each genotype. (F) Quantification of cell number per cluster for each genotype. *** p≤0.001 two-tailed unpaired t test. N=30 border cell clusters per genotype. (A-B, D-E) Scale bars 20μm. (A’-B’, D’-E’) Scale bars 5μm.

We next asked if the observed reduction in cell numbers within the border cell cluster was due to Rap1 function in border cell specification or in the recruitment of cells due to cell survival. We examined anterior follicle cells at stage 8 for a reporter of JAK/STAT activity, 10XSTAT::GFP (Bach et al., 2007). These cells are fated to become the migratory border cells (Beccari et al., 2002; Silver and Montell, 2001). When both polar cells were present, we did not detect changes in either the pattern of 10XSTAT:GFP or the levels of nuclear STAT, and hence activity, in the three follicle cells immediately adjacent to the polar cells at stages 7-8 in DN-Rap1^N17^ compared to control (Fig. S5A-C). These results suggest that border cell fate specification at stage 8 of oogenesis is normal under conditions when both polar cells are present and does not rely on Rap1 activity. However, we cannot rule out a subtle decrease in the JAK/STAT activity gradient when only one polar cell survives to stage 8 in DN-Rap1^N17^ egg chambers.

Finally, we determined if the smaller number of cells per border cell cluster was due to the increased apoptotic activity observed upon Rap1 inhibition. We co-expressed DN-Rap1^N17^ with either a UAS-LacZ control or the apoptosis inhibitor p35 (Fig. 7D-F). We quantified the number of cells per cluster at stages 9 and 10, when border cell clusters have already formed and have either begun to migrate or finished their migration. Border cell clusters that co-expressed DN-Rap1^N17^ and LacZ had an average of 4.8 cells per cluster, resembling the phenotypes observed when expressing DN-Rap1^N17^ alone (Fig. 7D, D’, F compared to Fig. 7C). In contrast, border cell clusters co-expressing DN-Rap1^N17^ and p35 had an average of 6.4 cells per cluster (Fig. 7E-F), matching control border cell clusters (Fig. 7A, A’, C). Thus, the observed smaller border cell clusters caused by loss of Rap1 activity is likely due to apoptotic cell death of mature polar cells just prior to recruitment of border cells. Together, these data support a model in which Rap1 promotes mature polar cell survival by preventing apoptosis through upregulation of DIAP1, thus promoting an optimal number of cells being recruited to form the migrating border cell cluster.

## Discussion

A key challenge is to understand how epithelia and constituent cells survive and maintain shape in response to the changes in tissue shape and size during normal development. Here we focused on understanding how the small GTPase Rap1 maintains cell and tissue shapes during *Drosophila* egg chamber growth. Prior work has shown requirements for Rap1 in epithelial morphogenesis (Asha et al., 1999; Boettner and Van Aelst, 2007; Choi et al., 2013; Bonello et al., 2018), but whether and how Rap1 maintains epithelia during tissue growth and homeostasis was unclear. Here we used a model of developmental tissue growth in the *Drosophila* ovary to interrogate the function of Rap1. We found that Rap1 promotes epithelial follicle cell shape, polar cell shape, and local tissue shapes during a period of major egg chamber growth. We propose that this function of Rap1 in maintenance of epithelial integrity is through regulation of dynamic linkages between adherens junctions and the contractile actomyosin cytoskeleton. Our experiments revealed an unexpected and new role for Rap1 in regulating cell survival, especially of the mature polar cells, which leads to the assembly of a migratory border cell cluster with the optimal number of cells.

### Rap1 maintains cell and tissue shapes by modulating adherens junction-cytoskeleton linkages during tissue growth

Here we report that Rap1 is required to maintain polar cell and follicle cell shapes and proper tissue shapes of the anterior follicular epithelium during egg chamber elongation. Our quantitative analyses of individual cell shapes and local tissue deformation demonstrated that Rap1 maintained tissue and epithelial morphology. Moreover, Rap1-deficient polar cell shapes were highly affected. Most of these phenotypes were at stages 7-8, a period of dramatic egg chamber growth (Crest et al., 2017). Previous work using *Rap1* null flies that were rescued to viable adults by expression of heat shock-driven Rap1 revealed egg chambers that degenerated and had distorted follicle cell shapes (Asha et al., 1999). Notably, these phenotypes seemed to be more severe during mid-to-late stages of oogenesis, stages that overlap with egg chamber elongation, thus supporting a role for Rap1 in tissue maintenance. We obtained similar, albeit less severe egg chamber defects, by specifically expressing DN-Rap1^N17^ in follicle cells. These results indicate a requirement of Rap1 in tissue shape maintenance.

How does Rap1 contribute to cell and tissue shape maintenance within the follicular epithelium during tissue expansion? Rap1 regulates early Bazooka/Par3 localization and adherens junction positioning during *Drosophila* embryogenesis (Bonello et al., 2018). *Rap1* mutant embryos fail to localize spot adherens junctions properly during cellularization and complete embryogenesis with fragmented cuticles suggesting a loss of tissue integrity (Choi et al., 2013). Similarly, Rap family proteins are required for adherens junctions and tight junction formation in MDCK cells (Sasaki et al., 2020). Our results showed that Rap1 is required specifically for the enrichment of E-Cadherin at the apical side of polar cells, which we propose helps promote proper polar cell shapes. We also observed stretching of the anterior epithelial follicle cells, which may indicate altered adhesion in these cells. Although E-cadherin appeared to be localized to apical puncta between follicle cells consistent with the formation of adherens junctions, these junctions may not be completely normal thus contributing to the stretched epithelial shapes. We thus propose that Rap1 maintains (or helps assemble) the proper positioning of the apical adherens junction, at least in polar cells. Further work will be needed to determine if and how Rap1 mechanistically maintains adherens junctions in the follicle cells.

The functions of adherens junctions to maintain cell-to-cell contacts within an epithelium are also coupled to actomyosin contractility, which can drive cell shape changes. Rap1 and its effector Canoe (Cno) regulate actomyosin contractility in apically constricting mesodermal cells during *Drosophila* ventral furrow invagination (Sawyer et al., 2009; Spahn et al., 2012). Similarly, Rap1 is required for Shroom dependent apical constriction in *Xenopus* (Haigo et al., 2003). These studies led us to ask if, in addition to its role in positioning adherens junctions during egg chamber elongation, Rap1 was responsible for proper actomyosin contractility. We found that Myo-II was required for proper tissue shape during egg chamber elongation, but surprisingly Rap1 was dispensable for Myo-II localization in polar cells. Taken together with the stretched individual cell shapes and deformed egg chambers observed for α-Catenin RNAi, these data support our model that Rap1 maintains the strength of adherens junction-actin linkages as was reported for the known Rap1 effector Cno (Sawyer et al., 2009).

We found that Rap1 regulates egg chamber shape during a period of major tissue growth. The inherent on-off activity states of GTPases like Rap1 make them particularly useful for dynamic processes that require discrete bursts of activity in response to cellular or tissue level cues (Gloerich and Bos, 2011). In apical constriction, for example, myosin pulsatility is coupled to progressive tightening of the apical domain (Martin and Goldstein, 2014; Martin et al., 2009). The transient nature of pulsatile myosin may require a fast-acting molecular switch that can be activated and inactivated quickly to couple motor behaviors to changes at the cell cortex. While we did not observe Myo-II localization defects when Rap1 was inhibited, the requirement for α-Catenin indicates that linkage to the actomyosin cytoskeleton is critical for epithelial cell and tissue shapes. Atypical Protein Kinase C (aPKC) promotes an optimal level of actomyosin to maintain an intact and organized follicle cell epithelium (Osswald et al., 2022). Indeed, acute loss of aPKC causes the follicle cell layer to apically constrict and rupture, due to growth of the egg chamber. While the effects caused by loss of Rap1 activity did not cause the epithelium to rupture, the tissue instead locally deformed. The basement membrane becomes thinner at the anterior and posterior ends during tissue elongation, allowing further growth along the anterior-posterior axis (Balaji et al., 2019; Crest et al., 2017). We suggest that loss of Rap1, in conjunction with a permissive region of the basement membrane, allows the tissue to widen in response to egg chamber growth. Thus, we propose that Rap1, through apical enrichment of adherens junction proteins, reinforces strong epithelial connections and resistance to growth of the germline.

The signal relay mechanisms that act upstream to regulate the various Rap1 activities in the follicle cell epithelium are unknown. Nor is it known which Rap1 effectors mediate the direct control of adherens junction-actomyosin linkages in this context. The GTPase activating protein Rapgap1 acts as one regulatory layer controlling Rap1 α-Catenin modulation in regulating dorsal fold morphogenesis and is expressed in the ovary (Wang et al., 2013; Sawant et al., 2018). *Drosophila* has several known guanine nucleotide exchange factors (GEFs), including PDZ-GEF (also known as Dizzy), C3G, and Epac. Whether one or more GEFs have differential functions during these stages of oogenesis, and in cell survival, remains to be tested. A well-known Rap1 effector, Cno, acts as a key signal relay mechanism downstream of Rap1. For example Cno mediates Rap1 functions in *Drosophila* morphogenesis including mesoderm invagination, head involution, and dorsal closure through modular biochemical functions (Perez-Vale et al., 2021; Sawyer et al., 2009). Whether these or other upstream and downstream regulators mediate Rap1 functions in epithelial tissue maintenance and cell survival remain to be determined.

### Rap1 promotes follicle cell and polar cell survival during oogenesis

We found that Rap1 maintains epithelial cell viability during oogenesis. One possibility is that the role for Rap1 in adherens junction protein localization is coupled to cell survival. Cell-cell contacts are essential for cell viability in certain contexts (Guillot and Lecuit, 2013). We found that the adherens junction-cytoskeleton linker protein α-Catenin not only supported proper epithelial and tissue shapes, but also promoted cell survival during oogenesis. Thus, cell-cell adhesions may be coupled to cell viability during tissue growth during oogenesis. Indeed, this period of dramatic tissue growth places extra strain on the epithelial follicle cells (Osswald et al., 2022), which could result in fewer cells surviving. Another major role for Rap1 was in promoting survival of the mature polar cells. The consequence of fewer polar cells surviving was a smaller border cell cluster, which is critical for optimal migration speed and ability to reach their final position at the oocyte (Cai et al., 2016; Stonko et al., 2015). Thus, Rap1’s role in promoting cell viability is critical for normal oogenesis.

We do not yet have a clear understanding of the mechanism by which Rap1 promotes cell survival and suppresses apoptosis. Our results suggest that Rap1 may have independent functions in cell survival other than (or in addition to) regulation of adherens junction proteins. Although knocking down α-Catenin resulted in apoptotic cells, the phenotype was overall much milder than that observed with Rap1 inhibition. Normally, DIAP1 must be maintained in the two mature polar cells to prevent their apoptosis (Borensztejn et al., 2013; Khammari et al., 2011). We observed a decrease in DIAP1 accumulation in the mature polar cells upon Rap1 inhibition, which is unlikely to be directly associated with defects in cell-cell adhesion. During stages 7 to 8, the overall levels of DIAP1 undergo a global reduction, which serves as a checkpoint mechanism to terminate unhealthy egg chambers rather than commit additional nutritional and energy resources (Baum et al., 2007). Rap1 promotes levels of DIAP1 before and during these stages, thus protecting the mature polar cells and likely other follicle cells from undergoing abnormal cell death. It remains to be tested whether Rap1 generally maintains cell survival of follicle cells by more directly fine tuning DIAP1 levels at the molecular level, or if the regulation of DIAP1 by Rap1 is indirect, for example due to disruption of epithelial integrity when Rap1 is inhibited. It is also unclear how cellular mechanics function together with transcription of DIAP1 to promote cell survival during tissue growth of the ovary. Further work will also be needed to determine if the function for Rap1 in cell survival and maintenance of epithelial shapes in growing tissues is conserved in other developing tissues and organs.

## MATERIALS AND METHODS

### *Drosophila* genetics

All fly stocks used in this study are listed in Table 1 and the complete genotypes for each experiment can be found in Table 2. Crosses were typically set up and maintained at 25°C. In cases where transgene expression impacted organism viability, the crosses were set up and maintained at 18°C. The tub-GAL80ts (‘tsGAL80’) transgene (McGuire et al., 2004) was present in the genetic background of many crosses in this study to repress GAL4 expression during other stages of development. Flies were shifted to 29°C for 12-72 hours prior to dissection to ensure optimal GAL4 expression and repression of tsGAL80, unless otherwise noted.

**Table 1.**
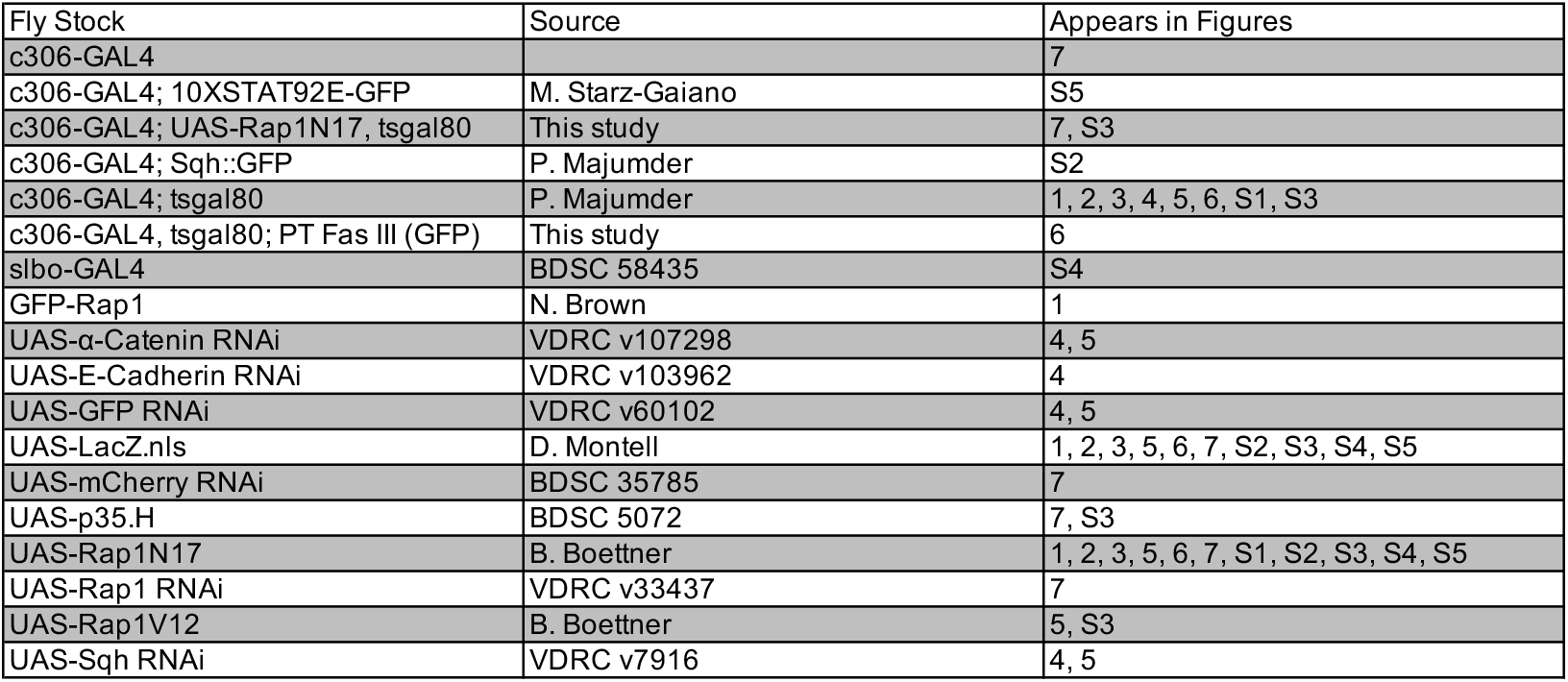
Fly strains used in this study.

| Fly Stock | Source | Appears in Figures |
| --- | --- | --- |
| c306-GAL4 |  | 7 |
| c306-GAL4; 10XSTAT92E-GFP | M. Starz-Gaiano | S5 |
| c306-GAL4; UAS-Rap1N17, tsgal80 | This study | 7, S3 |
| c306-GAL4; Sqh::GFP | P. Majumder | S2 |
| c306-GAL4; tsgal80 | P. Majumder | 1, 2, 3, 4, 5, 6, S1, S3 |
| c306-GAL4, tsgal80; PT Fas III (GFP) | This study | 6 |
| slbo-GAL4 | BDSC 58435 | S4 |
| GFP-Rap1 | N. Brown | 1 |
| UAS-α-Catenin RNAi | VDRC v107298 | 4, 5 |
| UAS-E-Cadherin RNAi | VDRC v103962 | 4 |
| UAS-GFP RNAi | VDRC v60102 | 4, 5 |
| UAS-LacZ.nls | D. Montell | 1, 2, 3, 5, 6, 7, S2, S3, S4, S5 |
| UAS-mCherry RNAi | BDSC 35785 | 7 |
| UAS-p35.H | BDSC 5072 | 7, S3 |
| UAS-Rap1N17 | B. Boettner | 1, 2, 3, 5, 6, 7, S1, S2, S3, S4, S5 |
| UAS-Rap1 RNAi | VDRC v33437 | 7 |
| UAS-Rap1V12 | B. Boettner | 5, S3 |
| UAS-Sqh RNAi | VDRC v7916 | 4, 5 |

**Table 2.** Genotypes in this study.

| Figure | Genotypes |
| --- | --- |
| 1A | w; P[w+ GFP-Rap1] |
| 1B | c306-GAL4/w; tsgal80/+; UAS-LacZ.nls/+ |
| 1C | c306-GAL4/w; tsgal80/+; UAS-LacZ.nls/+ |
| 1D | c306-GAL4/w; tsgal80/UAS-Rap1N17; TM2/+ |
| 1E | c306-GAL4/w; tsgal80/+; UAS-LacZ.nls/+ |
| 1F | c306-GAL4/w; tsgal80/UAS-Rap1N17; TM2/+ |
| 1G | c306-GAL4/w; tsgal80/+; UAS-LacZ.nls/+<br>c306-GAL4/w; tsgal80/UAS-Rap1N17; TM2/+ |
| 2A-A" | c306-GAL4/w; tsgal80/+; UAS-LacZ.nls/+ |
| 2B-B" | c306-GAL4/w; tsgal80/UAS-Rap1N17; TM2/+ |
| 2C | c306-GAL4/w; tsgal80/+; UAS-LacZ.nls/+<br>c306-GAL4/w; tsgal80/UAS-Rap1N17; TM2/+ |
| 3A-A" | c306-GAL4/w; tsgal80/+; UAS-LacZ.nls/+ |
| 3B-B" | c306-GAL4/w; tsgal80/UAS-Rap1N17; TM2/+ |
| 3C-D | c306-GAL4/w; tsgal80/+; UAS-LacZ.nls/+<br>c306-GAL4/w; tsgal80/UAS-Rap1N17; TM2/+ |
| 4A | c306-GAL4/w; tsgal80/UAS-GFP-RNAi |
| 4B | c306-GAL4/w; tsgal80/UAS-Sqh-RNAi |
| 4C | c306-GAL4/w; tsgal80/UAS-E-Cadherin-RNAi |
| 4D | c306-GAL4/w; tsgal80/UAS-α-Catenin-RNAi |
| 4E | c306-GAL4/w; tsgal80/UAS-GFP-RNAi<br>c306-GAL4/w; tsgal80/UAS-Sqh-RNAi<br>c306-GAL4/w; tsgal80/UAS-E-Cadherin-RNAi<br>c306-GAL4/w; tsgal80/UAS-α-Catenin-RNAi |
| 4F-F' | c306-GAL4/w; tsgal80/UAS-GFP-RNAi |
| 4G-G' | c306-GAL4/w; tsgal80/UAS-Sqh-RNAi |
| 4H-H' | c306-GAL4/w; tsgal80/UAS-α-Catenin-RNAi |
| 4I-I' | c306-GAL4/w; tsgal80/UAS-α-Catenin-RNAi |
| 5A | c306-GAL4/w; tsgal80/+; UAS-LacZ.nls/+ |
| 5B | c306-GAL4/w; tsgal80/UAS-Rap1N17; TM2/+ |
| 5C | c306-GAL4/w; tsgal80/+; UAS-LacZ.nls/+<br>c306-GAL4/w; tsgal80/UAS-Rap1N17; TM2/+<br>c306-GAL4/w; tsgal80/BL; UAS-Rap1V12/+ |
| 5D | c306-GAL4/w; tsgal80/UAS-GFP-RNAi |
| 5E | c306-GAL4/w; tsgal80/UAS-Sqh-RNAi |
| 5F | c306-GAL4/w; tsgal80/UAS-α-Catenin-RNAi |
| 5G | c306-GAL4/w; tsgal80/UAS-GFP-RNAi<br>c306-GAL4/w; tsgal80/UAS-Sqh-RNAi<br>c306-GAL4/w; tsgal80/UAS-α-Catenin-RNAi |
| 6A-A" | c306-GAL4/w; tsgal80/+; UAS-LacZ.nls/+ |
| 6B-B" | c306-GAL4/w; tsgal80/UAS-Rap1N17; TM2/+ |
| 6C-C" | c306-GAL4, tsgal80/w; PT Fas III (GFP)/+; UAS-LacZ.nls/+ |
| 6D-D" | c306-GAL4, tsgal80/w; PT Fas III (GFP)/+; UAS-LacZ.nls/+ |
| 6E-E" | c306-GAL4, tsgal80/w; PT Fas III (GFP)/UAS-Rap1N17; TM2/+ |
| 6F-F" | c306-GAL4, tsgal80/w; PT Fas III (GFP)/UAS-Rap1N17; TM2/+ |
| 7A-A' | c306-GAL4/w; G13, L[2]/+; UAS-LacZ.nls/+ |
| 7B-B' | c306-GAL4/w; G13, L[2]/UAS-Rap1N17; TM2/+ |
| 7C | c306-GAL4/w; G13, L[2]/+; UAS-LacZ.nls/+ |
|  | c306-GAL4/w; G13, L[2]/UAS-Rap1N17; TM2/+ |
|  | c306-GAL4/w; G13, L[2]/+; UAS-mCherry-RNAi/+ |
|  | c306-GAL4/w; G13, L[2]/+; UAS-Rap1-RNAi/+ |
| 7D-D' | c306-GAL4/w; tsgal80, UAS-Rap1N17/+; UAS-LacZ.nls/+ |
| 7E-E' | c306-GAL4/w; tsgal80, UAS-Rap1N17/UAS-p35.H |
| 7F | c306-GAL4/w; tsgal80, UAS-Rap1N17/+; UAS-LacZ.nls/+ |
|  | c306-GAL4/w; tsgal80, UAS-Rap1N17/UAS-p35.H |
| S1A-B | c306-GAL4/w; tsgal80/UAS-Rap1N17; TM2/+ |
| S2A-A" | c306-GAL4/w; Sqh::GFP/+; UAS-LacZ.nls/+ |
| S2B-B" | c306-GAL4/w; Sqh::GFP/UAS-Rap1N17; TM2/+ |
| S2C | c306-GAL4/w; Sqh::GFP/+; UAS-LacZ.nls/+ |
|  | c306-GAL4/w; Sqh::GFP/UAS-Rap1N17; TM2/+ |
| S3A | c306-GAL4/w; tsgal80/+; UAS-LacZ.nls/+ |
|  | c306-GAL4/w; tsgal80/UAS-Rap1N17; TM2/+ |
|  | c306-GAL4/w; tsgal80/BL; UAS-Rap1V12/+ |
| S3B | c306-GAL4/w; tsgal80, UAS-Rap1N17/+; UAS-LacZ.nls/+ |
| S3C | c306-GAL4/w; tsgal80, UAS-Rap1N17/UAS-p35.H |
| S3D | c306-GAL4/w; tsgal80, UAS-Rap1N17/+; UAS-LacZ.nls/+ |
|  | c306-GAL4/w; tsgal80, UAS-Rap1N17/UAS-p35.H |
| S4A | w; slbo-GAL4/UAS-LacZ.nls |
| S4B-B' | w; slbo-GAL4/UAS-LacZ.nls |
| S4C-C' | w; UAS-Rap1N17/+; slbo-GAL4/TM2 |
| S4D | w; slbo-GAL4/UAS-LacZ.nls |
|  | w; UAS-Rap1N17/+; slbo-GAL4/TM2 |
| S5A-A" | c306-GAL4/w; 10XSTAT92E-GFP/+; UAS-LacZ.nls/+ |
| S5B-B" | c306-GAL4/w; 10XSTAT92E-GFP/UAS-Rap1N17; TM2/+ |
| S5C | c306-GAL4/w; 10XSTAT92E-GFP/+; UAS-LacZ.nls/+ |
|  | c306-GAL4/w; 10XSTAT92E-GFP/UAS-Rap1N17; TM2/+ |

### Immunostaining

Antibodies, sources, and dilutions used are listed in Table 3. Fly ovaries from 2-to 8-day old females were dissected in Schneider’s *Drosophila* Medium (Thermo Fisher Scientific, Waltham, MA, USA) supplemented with 10% fetal bovine serum (Seradigm FBS; VWR, Radnor, PA, USA). Ovaries were either kept whole or dissected further into ovarioles and fixed for 10 mins using 16% methanol-free formaldehyde (Polysciences, Inc., Warrington, PA, USA) diluted to a final concentration of 4% in 1X Phosphate Buffered Saline (PBS). Following fixation, tissues were washed ≥4x with ‘NP40 block’ (50 mM Tris-HCL, pH 7.4, 150mM NaCl, 0.5% NP40, 5mg/ml bovine serum albumin [BSA]) and rocked in the solution for ≥30 mins prior to antibody incubation. Primary and secondary antibody incubation as well as all other subsequent wash steps were also performed in NP40 block. Dissected and stained ovarioles and egg chambers were mounted on slides with Aqua-PolyMount (Polysciences, Inc.) media and allowed to harden prior to imaging.

**Table 3.**
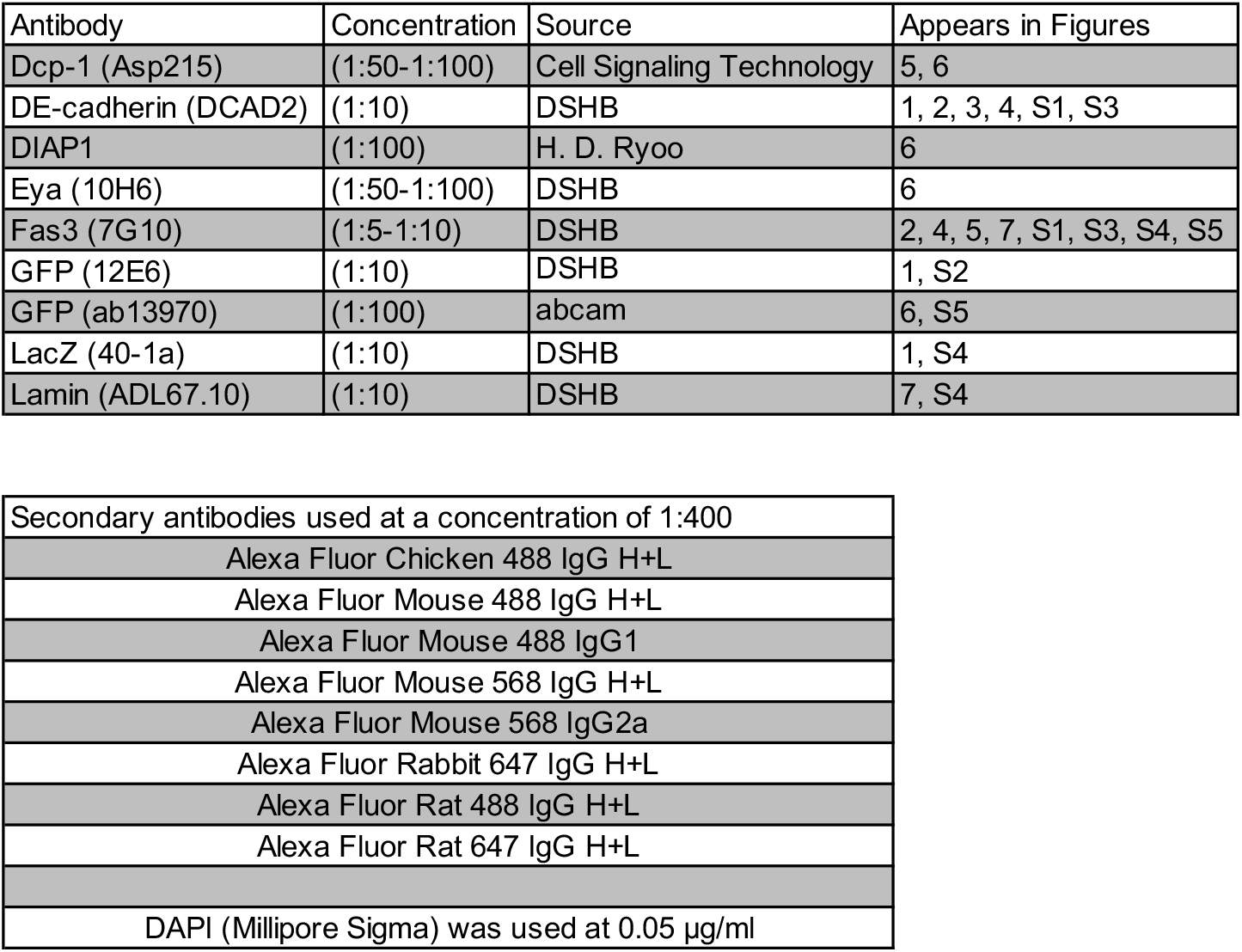
Antibodies used in this study.

| Antibody | Concentration | Source | Appears in Figures |
| --- | --- | --- | --- |
| Dcp-1 (Asp215) | (1:50-1:100) | Cell Signaling Technology | 5, 6 |
| DE-cadherin (DCAD2) | (1:10) | DSHB | 1, 2, 3, 4, S1, S3 |
| DIAP1 | (1:100) | H. D. Ryoo | 6 |
| Eya (10H6) | (1:50-1:100) | DSHB | 6 |
| Fas3 (7G10) | (1:5-1:10) | DSHB | 2, 4, 5, 7, S1, S3, S4, S5 |
| GFP (12E6) | (1:10) | DSHB | 1, S2 |
| GFP (ab13970) | (1:100) | abcam | 6, S5 |
| LacZ (40-1a) | (1:10) | DSHB | 1, S4 |
| Lamin (ADL67.10) | (1:10) | DSHB | 7, S4 |

|  |
| --- |
| Secondary antibodies used at a concentration of 1:400 |
| Alexa Fluor Chicken 488 IgG H+L |
| Alexa Fluor Mouse 488 IgG H+L |
| Alexa Fluor Mouse 488 IgG1 |
| Alexa Fluor Mouse 568 IgG H+L |
| Alexa Fluor Mouse 568 IgG2a |
| Alexa Fluor Rabbit 647 IgG H+L |
| Alexa Fluor Rat 488 IgG H+L |
| Alexa Fluor Rat 647 IgG H+L |
| DAPI (Millipore Sigma) was used at 0.05 µg/ml |

### Microscopy

Images of fixed egg chambers were acquired on a Zeiss LSM 880 confocal laser scanning microscope (KSU College of Veterinary Medicine Confocal Core) using either a 20 × 0.75 numerical aperture (NA) or a 40 × 1.3 NA oil-immersion objective controlled by Zeiss Zen 14 software.

### Image Processing and Data Analysis

Measurements were performed using FIJI (Schindelin et al., 2012). Egg chamber local deformation measurements were determined by measuring the width between the apical follicle cell surfaces at 20% of the anterior-posterior (A-P) length of the egg chamber (the very anterior tip is 0% of the A-P length). The polar cell aspect ratio was measured by analyzing Z-stacks taken of egg chambers stained with FasIII (to identify polar cells), E-Cadherin, and DAPI; the length and width was measured at the widest point of each polar cell. The aspect ratio was then calculated by dividing the width by length.

E-Cadherin accumulation in polar cells (identified by FasIII) was measured by analyzing Z-stacks acquired of egg chambers stained for E-Cadherin, FasIII, and DAPI. A 7μm line was drawn to quantify pixel intensity, starting within the germline (an adjacent nurse cell), extending through the apical polar cell-polar cell contact, then continuing along the lateral interface (see line in Fig. 3A’ and 3B’). Only images in which both polar cells could be viewed in the same Z-slice were used for quantitation. The “Plot Profile” function in FIJI was used to obtain a list of pixel intensity values corresponding to points along this line. These values were normalized to the highest pixel intensity measured in the experiment and plotted. Staining and imaging conditions were kept consistent between samples.

Dying cells during oogenesis were quantified by scanning through dissected whole ovarioles stained and imaged for cDcp-1, DAPI, and FasIII. Nuclei positive for caspase activity (cDcp-1-expressing nuclei) from stages 2 through 8 were quantified. Death of mature polar was scored by analyzing cDcp-1 expressing nuclei specifically in stage 7-8 egg chambers. DIAP1 accumulation in polar cells was assessed by acquiring Z-stacks through the polar cells in stages 4-6 and stages 7-8 egg chambers and analyzing qualitative reduction in the DIAP1 signal compared to adjacent cells. Polar cells were identified using a GFP protein trap in FasIII.

Quantification of the number of cells per border cell cluster were measured by acquiring Z-stacks through border cell clusters visualized using the nuclear envelope marker Lamin (also known as Lamin Dm_0_) and DAPI. Whenever possible, E-Cadherin was used to determine the boundaries of fully delaminated border cell clusters. Z-stacks encompassing the entire cluster including both border cells and polar cells were acquired and nuclei were manually counted using the Lamin signal.

Apical Myo-II accumulation was quantified by acquiring Z-stacks of stage 7-8 egg chambers expressing a fluorescent Myo-II reporter, Sqh::GFP, and co-stained for E-Cadherin and DAPI. A 2μm line was drawn at the apical surface of polar cells (identified by enrichment of E-Cadherin) and the “Plot Profile” function in FIJI was used to obtain a list of intensity values for each line. The intensity values were normalized to the highest pixel intensity measured in the experiment and plotted. Samples of inferior staining quality were eliminated from analysis. Staining and imaging conditions were kept consistent between samples.

Supernumerary polar cell load during oogenesis was scored by scanning through whole ovarioles and counting the number of polar cells present at each pole using FasIII and Eya as markers for polar cell fate. Nuclear STAT intensity at stages 7-8 was quantified by measuring 10XSTAT::GFP reporter intensity by drawing lines across three nuclei on either side of polar cells The mean GFP intensity of each line was normalized to the mean DAPI signal for each nuclear measurement. All measurements were then normalized to the highest relative GFP intensity value measured in the experiment.

## Figures, graphs, and statistics

Images were processed in FIJI and figures were assembled using Affinity Photo (Serif, Nottingham, United Kingdom). Illustrations were designed in Affinity Photo. Graphs and statistical tests were performed using GraphPad Prism 7 or Prism 9 (GraphPad Software, San Diego, CA, USA). All statistical tests and significance levels are listed in the figure legends for the figures in which they appear and in Table 4.

**Table 4.** Statistics in this study.

| Figure | Experimental Unit | Measurement | Experiments per Genotype (n) | Statistical Test | Significance |
| --- | --- | --- | --- | --- | --- |
| 1G | Egg chamber | Anterior width | ≥17 | Unpaired two-tailed t test | ** |
| 2C | Polar cell | Aspect ratio | ≥26 | Unpaired two-tailed t test | ns, **** |
| 3C | Egg chamber | Apical E-Cad Intensity | ≥10 | N/A | N/A |
| 4E | Egg chamber | Anterior width | ≥19 | One-way ANOVA with Dunnett's Multiple Comparisons test | ns, ** |
| 5C | Ovariole | cDcp+ nuclei | ≥29 | One-way ANOVA with Dunnett's Multiple Comparisons test | ns, **** |
| 5G | Ovariole | cDcp+ nuclei | ≥20 | One-way ANOVA with Dunnett's Multiple Comparisons test | ns, ** |
| 7C | Border cell clusters | Cells per cluster | ≥26 | Unpaired two-tailed t test | * **** |
| 7F | Border cell clusters | Cells per cluster | 30 | Unpaired two-tailed t test | *** |
| S2C | Egg chamber | Apical Sqh Intensity | ≥8 | N/A | N/A |
| S3A | Egg chamber pole | Supernumerary polar cells | ≥30 | Fisher's exact test | ns, * |
| S3D | Polar cell | Aspect ratio | ≥34 | Unpaired two-tailed t test | ns |
| S4D | Border cell clusters | Cells per cluster | ≥25 | Unpaired two-tailed t test | **** |
| S5C | Nucleus | Stat reporter intensity | ≥12 | Unpaired two-tailed t test | ns |

Significance values: \* $P \leq 0.05$ , \*\* $P \leq 0.01$ , \*\*\* $P \leq 0.001$ , \*\*\*\* $P < 0.0001$ .

## Acknowledgements

We would like to thank Denise Montell, Michelle Starz-Gaiano, Nick Brown, and Hyung Don Ryoo, the Vienna *Drosophila* Resource Center (VDRC), the Bloomington *Drosophila* Stock Center (BDSC), and the Developmental Studies Hybridoma Bank (DSHB) at the University of Iowa for fly protocols, fly stocks, and antibodies. We also thank Gibson Hoefgen and Manuel Garcia for general project assistance and maintenance of *Drosophila* strains and Emily Burghardt, Rehan Khan, and Yujun Chen for helpful comments on the manuscript. We acknowledge the Confocal Core, funded by the Kansas State University (KSU) College of Veterinary Medicine, which provided use of the Zeiss LSM 880 confocal microscope. We thank the KSU Statistics Consulting Laboratory for statistics advice. This work was supported by the National Science Foundation (NSF 2027617) to J.A.M. and a KSU Johnson Cancer Research Center Graduate Student Summer Stipend Award to C.L.M. and J.A.M. The authors declare no competing financial interests.

## Author Contributions

C. Luke Messer: Conceptualization, Formal analysis, Validation, Investigation, Visualization, Methodology, Writing - Original Draft, Writing - Review & Editing; Jocelyn A. McDonald: Conceptualization, Supervision, Funding acquisition, Methodology, Writing - Original Draft, Writing - Review & Editing.

---

**Figure S1.**
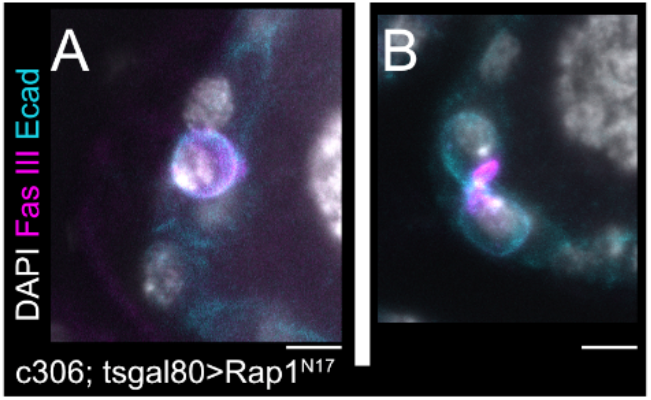
Rap1 inhibition causes polar cell loss and distorted polar cell shapes. (A-B) Anterior regions of two different DN-Rap1^N17^ stage 7-8 egg chambers. (A) Only one mature polar cell is present. (B) A pair of polar cells stretched along the dorsoventral axis. Scale bars 5μm.

**Figure S2.**
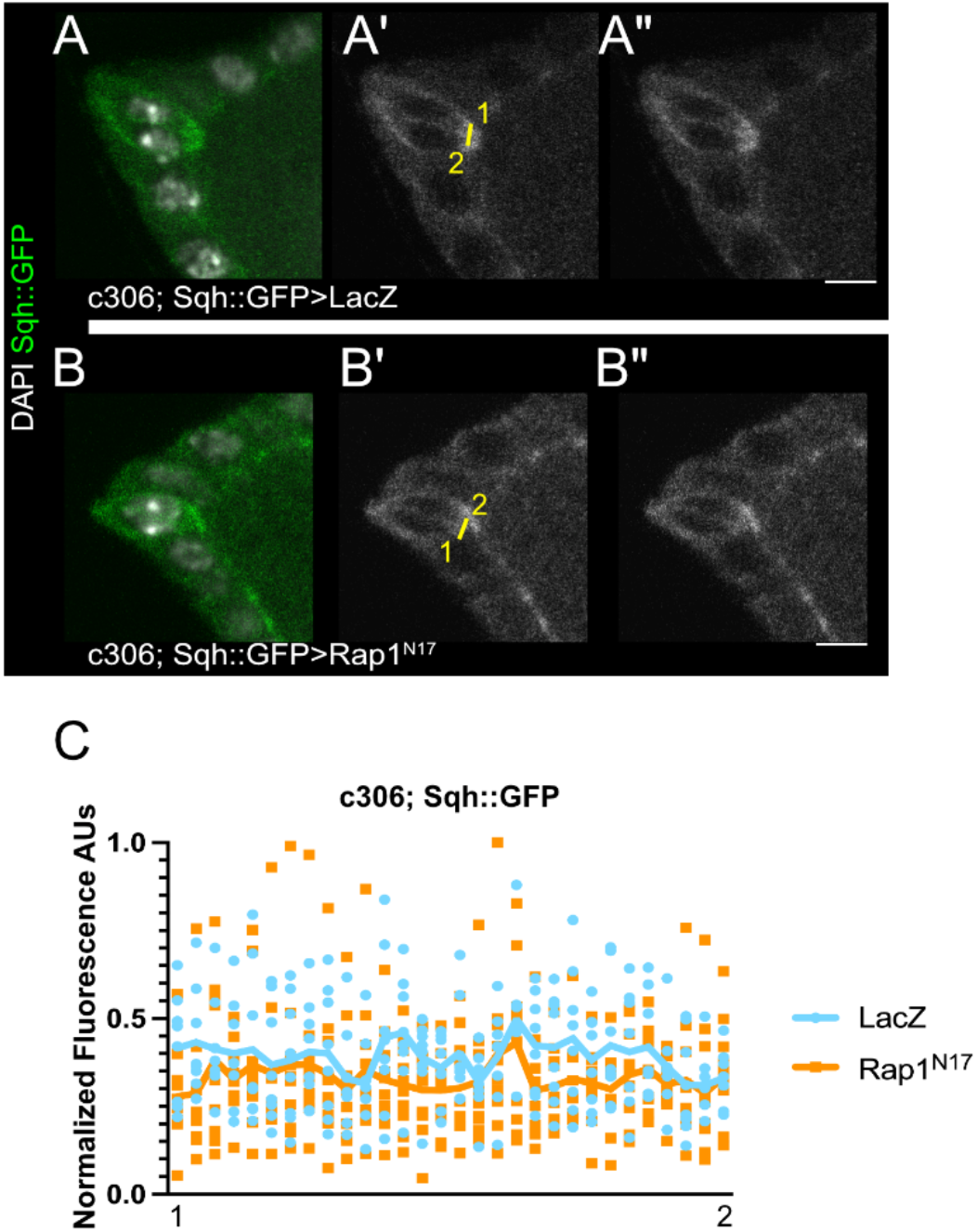
Rap1 is dispensable for Sqh apical localization in polar cells. (A-B) Anterior regions of stage 7-8 egg chambers for LacZ (A) or DN-Rap1^N17^ (B). (A’-B’) Single channel images of A and B showing GFP (gray) with yellow lines indicating region of measurement for Sqh::GFP intensity. The numbers correspond to the plot intensity profiles in C. (A”-B”) Single channel images as in A’ and B’ with line removed for clarity. The image brightness was adjusted for presentation purposes but not used for quantification. (C) Profile of the intensity values along the measurement lines, plotted and normalized to highest signal. The numbers on the x-axis correspond to the lines drawn on A’ and B’. Solid plot lines represent the mean intensity. N ≥ 8 egg chambers per genotype. Scale bars 5μm.

**Figure S3.**
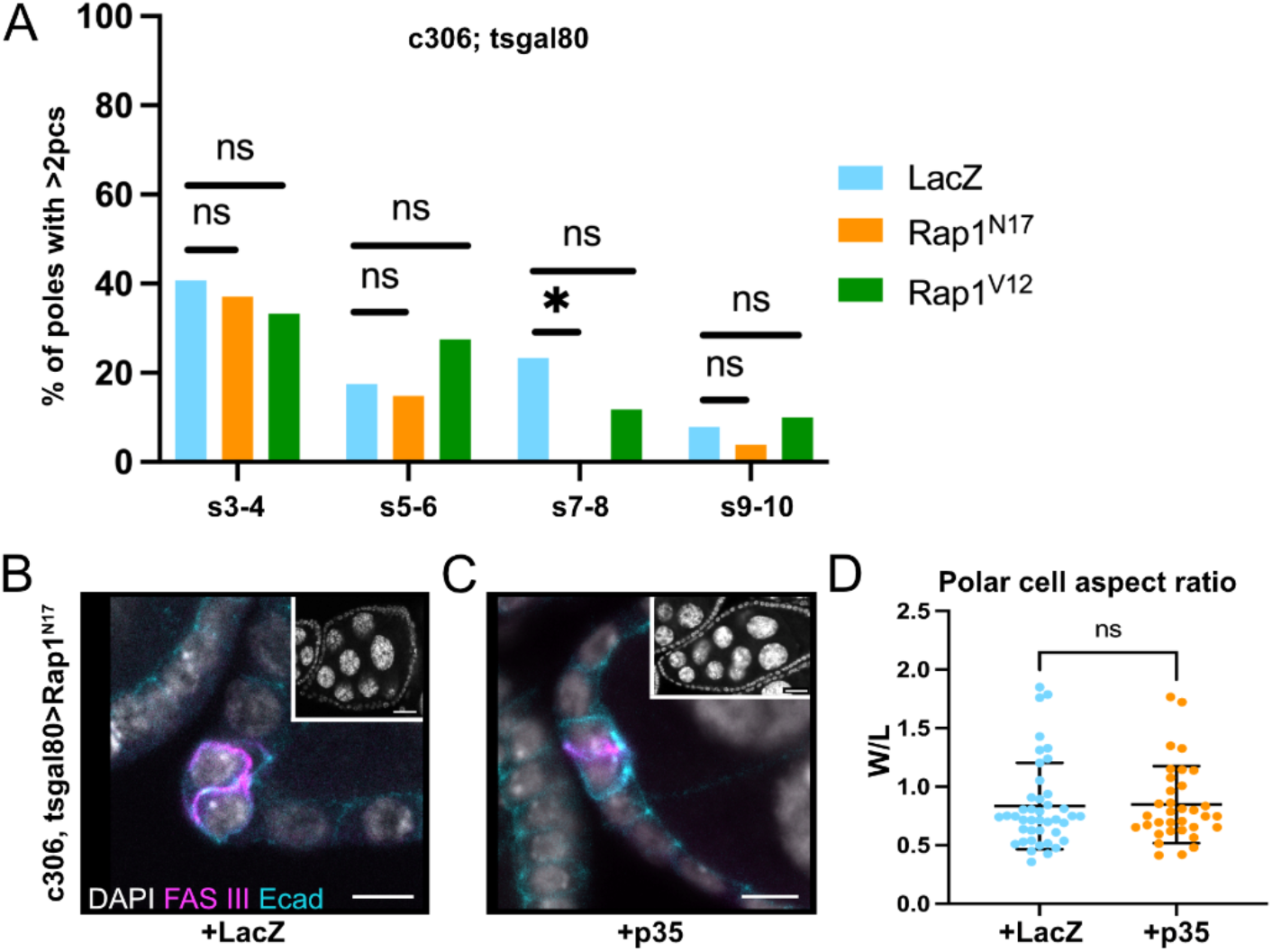
Rap1 does not regulate early elimination of supernumerary polar cells and maintains mature polar cell shape independent of apoptosis. (A) Quantification showing frequency of egg chamber poles with supernumerary polar cells during specific stages of oogenesis. * p≤0.05 two-sided Fisher’s exact test. N≥30 egg chamber poles per stage per genotype. Supernumerary polar cells are present early for all genotypes. (B-C) Anterior region of stage 7-8 egg chambers, DN-Rap1^N17^ with LacZ expression (A) or DN-Rap1^N17^ with apoptosis inhibitor p35 (B). Whole egg chambers stained for DAPI are shown in the insets. (D) Polar cell aspect ratio quantification. Mean AR of DN-Rap1^N17^ + LacZ control = 0.84. Mean AR of DN-Rap1^N17^ + p35 = 0.85. NS two-tailed unpaired t test. N≥34 polar cells per genotype. Main panel scale bars 5μm. Inset scale bars 20μm.

**Figure S4.**
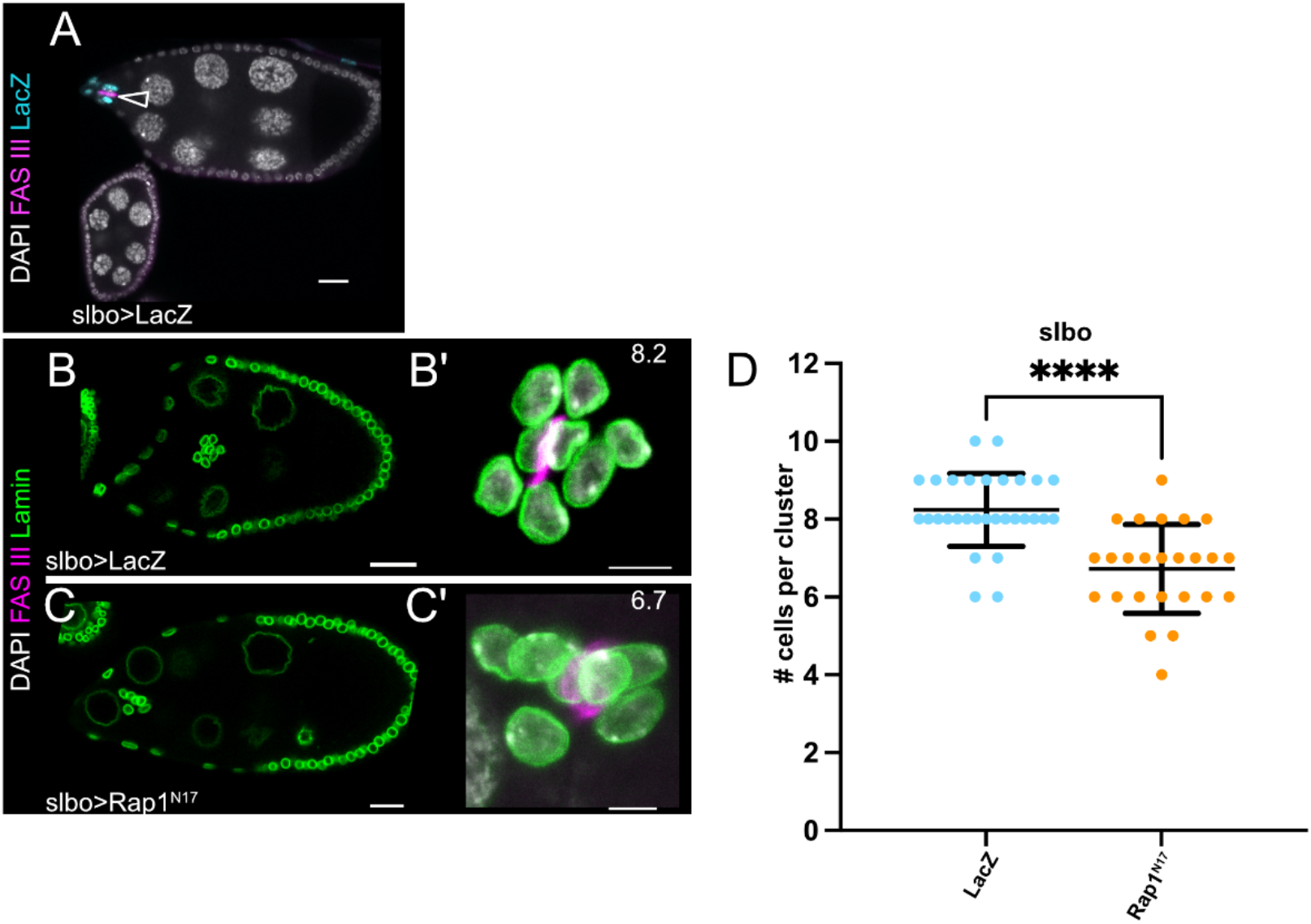
Effects of late Rap1 inhibition on border cell numbers. (A) Ovariole region expressing UAS-LacZ indicates the expression pattern of *slbo*-GAL4. (B-C) Stage 9-10 LacZ (B) and DN-Rap1^N17^ (C) egg chambers showing mid-migration border cells. (B’-C’) Maximum intensity projections of border cell clusters from B and C. Numbers in the upper right indicate the average number of cells per cluster. (D) Quantification of cell number per cluster for Rap1 inhibition using *slbo*-GAL4. **** p≤0.0001 two-tailed unpaired t test. N≥25 border cell clusters per genotype. (A-C) Scale bars 20μm. (B’-C’) Scale bars 5μm.

**Figure S5.**
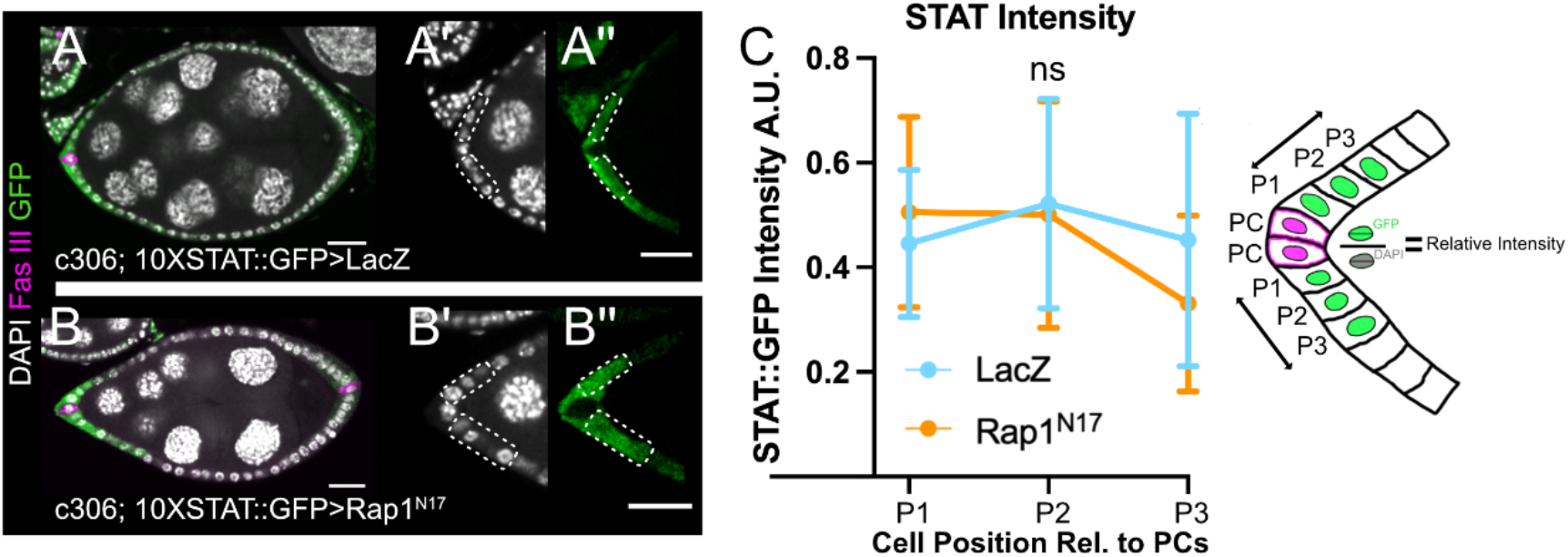
Rap1 is dispensable for STAT levels in follicle cells fated to become border cells. (A, B) Stage 8 LacZ (A) and DN-Rap1^N17^ (B) egg chambers with 10XSTAT::GFP reporter expression enriched adjacent to anterior polar cells. (A’-A”, B’-B”) Insets of the anterior end of the egg chambers in A and B, which was used to measure the intensity of nuclear 10XSTAT::GFP. (A’-B’) DAPI. (A”-B”) 10XSTAT::GFP. Dashed rectangles indicate the cell positions next to polar cells used for analyzing STAT levels. (C) Schematic and plot of STAT intensity values relative to the DAPI signal and normalized to the highest intensity STAT measurement. NS, unpaired two-tailed t-test. N≥12 follicle cells at each position pooled from N≥6 stage 8 egg chambers for each genotype. (A-B) Scale bars 20μm. (A”-B”) Scale bars 5μm.

## Notes

### Competing Interest Statement

The authors have declared no competing interest.

### Summary of Updates

Text has been updated, particularly in the Introduction.

